# New Δ246p53 isoform responds to DNA damage to enhance p53 family functions in cellular senescence

**DOI:** 10.1101/2023.04.07.536059

**Authors:** Shrutee N. Parkar, Ana Catarina Ramalho, Maria José López-Iniesta, Ricardo Silva, Jingyuan Zhao, Koto Kimura, Vanessa Dassi, Filipa da Silva Rita, Yari Ciribilli, Alessandra Bisio, Luísa Romão, Marco M. Candeias

## Abstract

*p53* is with little doubt one of the most powerful genes in our genome, as it makes growth *vs* arrest, repair *vs* replacement, catabolism *vs* anabolism, life *vs* death decisions in the cell. An alteration or malfunction in *p53* may lead to cancer, degeneration or premature ageing. So, it is not surprising that *p53* is also one of the most complex and tightly regulated genes, encoding for at least 10 RNA variants and 12 widely accepted protein forms. Here we identify one new p53 protein isoform of approximately 18 kDa that we termed Δ246p53. Δ246p53 is translated from an alternative translation initiation site (TIS) in codon 246. TIS-246 is preceded by a strong Kozak sequence and appears conserved in vertebrates, from sea lamprey to humans. Δ246p53’s origin and expression in cells were confirmed by frameshift and start codon mutations; five different antibodies against several epitopes from its N-terminus down to its C-terminus and one against the DNA binding domain region just upstream of its initiator methionine working as negative control; as well as siRNAs and an antisense oligo targeting TIS-246, which knocked-down Δ246p53 with little or no effect on full-length (FL) p53 protein levels. Δ246p53 was induced by DNA damage and triggered senescence and impaired tumour formation/growth in colony formation assays. Mechanistically, Δ246p53 interacted with FLp53, leading to a decrease in the expression of downstream target genes *HDM2* and *p21*. Concurrently, it also interacted with ΔNp63, leading to *p53*-independent *p63*-mediated activation of *p21*, a known senescence activator. Our results unveil a new naturally occurring and tightly controlled factor with specific functions in *p21* regulation and senescence. Future studies on Δ246p53 are likely to help us better understand and control the processes of tumour suppression and ageing.

## INTRODUCTION

By protecting the tissue – not the cell, as this one might be sacrificed to safeguard the health and functionality of the tissue as a whole – from DNA damage and other identity-compromising insults such as oncogene activation, infection or endoplasmic reticulum (ER) stress, *p53* manages development, repair, tumour suppression/progression and ageing. With this in mind, we may argue that *p53* first appears in organisms with true tissues, the Histozoa (Eumetazoa)^1^, because they are the first to be willing to sacrifice cell life for specialized tissue function and organismal fitness. Numerous motifs, residues, sequences, structures and domains in *p53* and its products are responsible for sensing the dangers threatening each tissue and activating the appropriate effectors/mechanisms to resolve each particular impasse. With the rising complexity of the organisms and increasing number of specialized tissues and cells, so evolved the complexity of *p53*. Human *p53* now produces, in different tissues and conditions, more than 10 different RNA and 12 different protein products^2–4^. Which isoform appeared first, and for what specific purpose, is still under debate. In general, shorter isoforms lacking both transactivation domains (TA) in the N-terminus (Δ133p53, Δ133p53β and Δ160p53) favour proliferation, survival and migration/invasion^5,6^; while forms maintaining at least one of the TA (full-length, FL; Δ40p53; p53β) promote apoptosis and cell-cycle arrest^7–9^. Things are less straightforward when it comes to DNA repair and ageing. FLp53 and Δ133p53 are both inducible by DNA damage and support DNA repair^7,10–12^; however, Δ160p53 is not induced by DNA damage unless integrated stress response (ISR) is also activated^13^. Regarding ageing, several p53 mouse models show accelerated ageing and most of them express an extra C-terminal portion of p53 (Δ40p53^14^ or the M protein^15^) or possess a mutated N-terminus^16,17^ (and thus an extra wild-type C-terminus compared to N-termini). So, while the N-terminus (or a complete p53 protein) seems to be associated with apoptosis, the C-terminus of p53 seems to be critical for DNA repair, senescence and ageing.

We have recently analysed the sequences of 50 different species from the sea vase to human and observed that the Δ160p53 protein product is likely to have appeared for the first time in mammals and its translation initiation site (TIS) is still conserved in most of them, while Δ133p53 only appeared later, in primates^18^. Here, we identified what could be one of the oldest and most conserved protein products of *p53*, dating back to Cambrian Stage 3 when vertebrates first emerged 500 million years ago. It is also the smallest natural p53 protein known to date, a “mini” p53 so to say, with only 148 amino acids and weighting around 18 kDa. Following the main trend for the nomenclature of p53 isoforms, we termed our protein Δ246p53 as it initiates at codon 246 of *p53*. Δ246p53 activated the *p21* gene in a *p63*-dependent manner and led to senescence and tumour suppression in the colony formation assay. It did not, however, in our experiments, induce cell death. In light of these findings and previous reports on the effect of C-terminal fragments of p53 in tumour resistance and premature ageing^15^, it will be interesting to investigate in the future how this naturally occurring isoform moulded the shape and lifespan of vertebrates and how it might be controlled to prolong life expectancy and improve well-being in old age.

## MATERIAL AND METHODS

### Cell culture and treatments

All cell lines were acquired from the American Type Culture Collection (ATCC), except for ASF-4-1 (registration number IFO50418) and ASF-4-5R1 (registration number JCRB1709), which were established by Kaji, K. and acquired from the Japanese Collection of Research Bioresources (JCRB). Cells were frequently tested for mycoplasma and other contaminations. The cell lines H1299, A549, HEK293T, LN229, SW-480 were maintained and cultured in Dulbecco’s Modified Eagle Medium (DMEM) (high glucose); U2OS, HCT116 and Saos2 were maintained and cultured in McCoy’s 5A (Modified) Medium; HEPG2, HT-1080, MCF-7, ASF-4-1, ASF-4-5R1 were maintained and cultured in Eagle’s Minimum Essential Medium (EMEM); with 10% fetal bovine serum. Media for all cells except ASF-4-1 and ASF-4-5R1 were also supplemented with 5 mM L-glutamine and Pencilin/Streptomycin 100X solution diluted 1:100. All cells were maintained at 37°C and 5% CO_2_. Thapsigargin (Sigma; 0.1 μM), Tunicamycin (Sigma; 1.2 μM), Etoposide (Sigma; 5 μM or as indicated), Oxaliplatin (Wako; 5 μM) and Paclitaxel (Funakoshi; 6 μM) treatments were for 16 (TH and TU) or 21 (ETO) or 24 (OXA and PTX) hours, or as indicated in the figure legends.

Transient transfections were performed according to the manufacturer’s protocol (293fectin, Invitrogen). To establish stable cell lines, pcDNA3.1 plasmids cloned with the desired inserts were linearized with PvuI (New England Biolabs) before transfection with 293fectin (Invitrogen) and two days later cells were selected using Neomycin (G418; Sigma; 1 mg/ml). Control non-transfected cells died several days after treatment and established cells showed similar levels of transgene expression as confirmed by Western blotting (WB). All p53 constructs were cloned into pcDNA3.1 using T4 DNA ligase (Takara).

### Western-blotting (WB) and immunoprecipitation (IP) assays

For WB, cells were lysed in 1.5x or 5x SDS sample buffer (62.5 mM Tris-HCl (pH 6.8), 12% glycerol, 2% SDS, 0.004% Bromo Phenol Blue, and 10% 2-mercaptoethanol) or NP40 lysis buffer (50mM Tris-HCl (pH 8.0) , 150mM NaCl, 1% Nonidet P40, 10% Glycerol) supplemented with Protease Inhibitor cocktail (PI, Roche, Merck/Sigma). After sonication, proteins were separated by SDS-PAGE on 10, 12, 14, or 15% gels and transferred to PVDF or Nitrocellulose membranes (Immobilon-FL, Millipore). After blocking for 1 h with blocking buffer (1X TBST with 5% w/v non-fat dry milk), membranes were blotted with antibodies diluted in blocking buffer, followed by incubation with secondary antibodies. Membranes were then soaked in Novex ECL (Thermo Fisher Scientific) or ECL Select (Cytiva, Amersham) or SuperSignal West Pico PLUS (Thermo Fisher Scientific) or SuperSignal West Atto (Thermo Fisher Scientific) and signals were captured with LAS-3000 Imager (Fujifilm) or Chemidoc MP Imaging System (BioRad).

For the immunoprecipitation assays, whole cell lysates were pre-cleared with mouse immunoglobulin G and protein G-sepharose before adding anti-HA antibody or DO-1 antibody. The beads were then washed 5 times in buffer A (1% Nonidet P-40, 150mM NaCl, 50mM Tris pH 8, 0.25% DOC, 1mM EDTA in the presence of Complete protease inhibitor cocktail (Roche)) and two times in PBS before boiling in 2x SDS sample buffer.

Most primary antibodies against p53 were gifts from Bořivoj (Borek) Vojtěšek (Masaryk Memorial Cancer Institute, RECAMO, Czech Republic): DO12, monoclonal (aa 256-267); Bp53.10, monoclonal (aa 374-378); Pab421, monoclonal (aa 376-378); CM1, polyclonal. KJC12, polyclonal, was a gift from Jean-Christophe Bourdon (University of Dundee, Dundee, United Kingdom). Other antibodies were as follows: anti-HA(Roche 3F10), anti-Lamin B1 (Santa Cruz Biotechnology A-11), anti-vinculin (Santa Cruz Biotechnology H-10 and 7F9), anti-p73 (Cell Signaling D3G10) monoclonal, anti-p63α Antibody (Selleck, F10N6), anti-p63 Antibody (Selleck, C16H5), anti-p73 Antibody (Selleck, F14N9). Secondary antibodies used were anti-mouse IgG HRP-linked antibody (Cell Signaling Technology, 1:3000), anti-rabbit IgG HRP-linked antibody (Cell Signaling Technology, 1:3000), anti-rat IgG HRP-linked antibody (Cell Signaling Technology, 1:3000) and anti-sheep IgG HRP-linked antibody (Jackson ImmunoLab, 1:6000).

### Knock-down assays

Morpholino antisense oligo was designed to target Δ246p53 translation initiation site and block translation initiation (MO; Gene Tools, OR, USA, 5’- GGTTCATGCCGCCCATGCAGGAACT -3’). Negative control Morpholino oligo used is a mutated version of MO (MO-ctl; Gene Tools, OR, USA; 5’-GG<u>A</u>TC<u>T</u>TGCCGC<u>G</u>CATGCA<u>C</u>GA<u>T</u>CT -3’). 5 μl of MO were added to the cells in 1 ml culturing media followed by 5 μl Endo-Porter delivery reagent and cells were harvested 3 days after. siRNA (GeneDesign) was delivered using Lipofectamine RNAiMAX (Life Technologies) or HiPerfect (Qiagen) following the manufacturers’ protocols. siRNA sequences for control and human *p53*, *p63*, *p73* and *SMAD3* are: Control (Ctl): 5’-CACCUAAUCCGUGGUUCAA-3’; p53-1 (ex7, s605 from Ambion); p53-2 (ex11, s607 from Ambion); p73(s14319, from Ambion): 5’-GCAAUAAUCUCGCAGUAtt-3’; p63(s229400, from Ambion) ; 5’-GAACCGCCGUCCAAUUUUAtt -3’; SMAD3(s8402, from Ambion): 5’-GUCUACCAGUUGACCCGAAtt -3’.

### MTT metabolic assay

MTT metabolic assay was performed according to the manufacturer’s protocol (MTT Cell Proliferation Assay Kit (Cayman Chemical, Ann Arbor, MI, USA)) using 4000 and 5000 cells treated or not with etoposide for 21 h. Calibration curves were also performed (2000 to 5000 cells).

### Senescence-associated β-galactosidase assay

Senescence-associated β-galactosidase assay was performed as previously published^19^. Briefly, cells were washed twice in PBS and then fixed with 4% paraformaldehyde for 5 min at room temperature and washed again three times in PBS. Freshly prepared SA-β-gal staining solution containing X-Gal was then added and cells incubated at 37°C in a humidified chamber for 16 h. Quantifications were performed automatically by Photoshop (Adobe), which calculates the percentage of surface covered in blue from 27 photographs taken randomly from a light microscope following a pre-established grid drawn under the culture wells and prepared in triplicate.

### Soft-agar colony formation assay

For soft-agar assays the experiments were carried out in twelve-well cell culture plates with Saos2 stable cells. The bottom layer corresponds to 500µl of 0.325% agar prepared with McCoys’s 5A Modified Medium (Gibco) supplemented with 15% (v/v) FBS, penicillin, streptomycin and 100 µg/mL G418. The upper layer of 0.6% agar containing 3000 cells per well was prepared by mixing equal volumes of a cell suspension with an agar solution and adding 400µl on top of the bottom agar layer. Single cell suspensions were prepared in 2x McCoys’s 5A Modified Medium, supplemented with 30% (v/v) FBS, 2% penicillin and streptomycin, and 200 µg/mL G418. After the medium solidified, they were overlaid with 200μl supplemented medium. The plates were incubated at 37°C in a 5% (v/v) CO_2_ atmosphere for 28 days and new medium was added every week. Colonies were stained with 0.1% crystal violet solution. The agar was washed with water before image acquisition for removal of excess crystal violet. Adobe Photoshop and Illustrator were used for standardized automated background removal and colony dot intensity enhancement using the same settings across all samples. Colony quantification was subsequently performed in ImageJ.

### Fluorescence-activated cell sorting (FACS)

Cells were harvested and fixed with ice-cold 70% ethanol. After 30-min incubation with RNase at 37°C, cells were stained with propidium iodide (PI, Sigma-Aldrich; 50 mg/ml) and cell cycle distribution was analysed using LSR Fortessa™ flow cytometer (BD Biosciences) and FlowJo Software [32]. The sub-G_0_/G_1_ population represents cells undergoing cell death.

### RNA isolation, RT and PCRs

Total RNAs were extracted from cultured cells using TRIzol reagent (Thermo Fisher Scientific) according to the manufacturer’s instructions. Complementary DNA was prepared using Superscript III reverse transcriptase (Thermo Fisher Scientific) with random hexamer primers according to the manufacturer’s instructions. The relative abundance of transcripts was finally evaluated by PCR. PCR primers used were as follows:

*p21* : 5’-CCTCAAATCGTCCAGCGACCTT-3’ and 5’-CATTGTGGGAGGAGCTGTGAAA-3’ *actin* : 5’-TCACCCACACTGTGCCCATCTACGA-3’ and 5’-TGAGGTAGTCAGTCAGGTCCCG-3’ qPCR was performed using Thunderbird^®^ SYBR™ qPCR Mix (cat. no.QPS-201, TOYOBO) according to the manufacturer’s instructions. 20 µL qPCR reaction mixes were prepared in 96-well plates and run on StepOnePlus Real Time PCR system (Thermo Fisher Scientific). No cDNA (skipping RT step) and water controls, as well as calibration curves, were performed in parallel.

Primers used for qPCR were as follows:

*p21* : 5’-TTCTGCTGTCTCTCCTCAGATTTCT-3’ and 5’-GGATTAGGGCTTCCTCTTGGA-3’ *HDM2* : 5’-CAGTAGCAGTGAATCTACAGGGA-3’ and 5’-CTGATCCAACCAATCACCTGAAT-3’ *GAPDH* (control housekeeping gene 1): 5’-CCATGAGAAGTATGACAACAGCC-3’ and 5’-GGGTGCTAAGCAGTTGGTG-3’ *RPS18* (control housekeeping gene 2): 5’-GCGGCGGAAAATAGCCTTTG-3’ and 5’-GATCACACGTTCCACCTCATC-3’

### Protein – protein interaction predictions

Predictions were performed using AlphaFold2-Multimer via the freely available online server^20^. FLp53 and Δ246p53 sequences were submitted as query sequence along with individual candidate binding partners : NF-κB(p50), NRF2, p63, p73, SMAD3 and SREBP1a. Predictions were performed using default settings with AlphaFold2-Multimer v2. Upon visualization the binding predictions were ranked based on AlphaFold’s predicted Local Distance Difference Test (pLDDT) score. The scores reflect the model’s confidence, which is colour coded based according to the following values: very high confidence (blue, pLDDT > 90), high confidence (cyan, 90 > pLDDT > 70), low confidence (yellow, 70 > pLDDT > 50), very low confidence (orange/red, pLDDT < 50).

### Sequence Analyses

Sequence alignment of *p53* from 50 different chordate species were performed using MAFFT^21^ server within Jalview. Likelihood of translation initiation from start codons homologous to human TIS-246 was analysed in 6 different mammalian species using NetStart 1.0^22^.

Sequences used are the following:

*Alligator sinensis* (Chinese alligator) XM_006038654.3

*Bos taurus* (cattle) NM_174201.2

*Callithrix jacchus* (marmoset) XM_002747948.4

*Callorhinchus milii* (elephant shark) JN794073.1

*Camelus bactrianus* (camel) XM_010965924.1

*Canis lupus familiaris* (dog) NM_001003210.1

*Carlito syrichta* (tarsier) XM_008062341.2

*Catharus ustulatus* (thrush) XM_033084255.1

*Cavia porcellus* (guinea pig) NM_001172740.1

*Ciona intestinalis* (vase tunicate) NM_001128898.1

*Danio rerio* (zebrafish) NM_001271820.1

*Dasypus novemcinctus* (armadillo) XM_012529094.2

*Dipodomys ordii* (kangaroo rat) XM_013013490.1

*Echinops telfairi* (lesser tenrec) XM_013007322.2

*Enhydra lutris kenyoni* (sea otter) XM_022524640.1

*Equus caballus* (horse) XM_023651624.1

*Erinaceus europaeus* (hedghog) XM_007523372.2

*Felis catus* (cat) NM_001009294.1

*Gallus gallus* (chicken) NM_205264.1

*Gorilla gorilla* (gorilla) XM_004058511.3

*Homo sapiens* (human) NM_001126112.3

*lctidomys tridecemlineatus* (squirrel) XM_005332819.3

*Latimeria chalumnae* (coelacanth) XM_005999799.2

*Loxodonta africana* (elephant) XM_010596586.2

*Macaca mulatta* (Rhesus monkey) NM_001047151.2

*Microcebus murinus* (mouse lemur) XM_012776058.2

*Mus musculus* (mouse) NM_011640.3

*Myotis lucifugus* (bat) XM_006102578.3

*Odobenus rosmarus* (walrus) XM_004398491.1

*Ornithorhynchus anatinus* (Platypus) ENSOANT00000053152.1

*Oryctolagus cuniculus* (rabbit) NM_001082404.1

*Oryzias latipes* (medaka) NM_001104742.1

*Otolemur garnettii* (galago) XM_012806041.2

*Pan troglodytes* (chimpanzee) XM_001172077.5

*Panthera pardus* (leopard) XM_019413568.1

*Pelodiscus sinensis* (turtle) XM_006112136.3

*Peromyscus leucopus* (white-footed mouse) XM_028869449.1

*Petromyzon marinus* (sea lamprey) XM_032962158.1

*Phascolarctos cinereus* (koala) XM_020966468.1

*Podarcis muralis* (lizard) XM_028752002.1

*Pongo abelii* (orangutan) XM_002826974.4

*Puma concolor* (puma) XM_025920288.1

*Rattus norvegicus* (rat) NM_030989.3

*Sarcophilus harrisii* (tasmanian devil) XM_031965445.1

*Sorex araneu* (shrew) XM_004604858.1

*Sus scrota* (pig) NM_213824.3

*Trichechus manatus* (manatee) XM_004376021.2

*Tursiops truncatus* (dolphin) XM_019944223.2

*Ursus arctos horribilis* (grizzly) XM_026520889.1

*Xenopus tropicalis* (frog) NM_001001903.1

## RESULTS

### 18 kDa band observed in cells expressing Δ133p53, Δ160p53 or mutant p53 is a translation product of *p53*

We have recently identified a new translation initiation site (TIS) in *p53* that produces a slightly smaller version of the Δ160p53 protein, Δ169p53, with similar pro-survival activity^18^. During our studies of these pro-oncogenic p53 forms in the *p53*-null human non-small cell lung carcinoma cell line H1299, we detected 4 distinct bands below Δ160/Δ169p53, ranging from around 26 kDa to approximately 18 kDa in size (**Fig. 1A, centre lane**). These were p53-specific bands as they were not observed in lysates of unmodified H1299 cells that lack the *p53* gene (**Fig. 1A, left lane**). The same bands were detected from Δ133p53 mRNA as well (**Fig. 1B**) though they were not readily visible from transient full-length (FL) p53 mRNA expression unless *p53* contained a mutation previously shown to activate the expression of short p53 isoforms^6^ (**Fig. 1C**). In order to discriminate if, or which of, these short p53 peptides are products of protein cleavage and/or products of alternative origins of protein synthesis, we inserted a frameshift mutation (one nucleotide deletion) in codon 157 (fs-157) upstream of the Kozak sequence for TIS-160 (translation initiation site for Δ160p53) in FL wild-type (WT) and mutant (R248Q) p53 transcripts carrying tag sequences for FLAG (at the 5’ end) and HA (at the 3’ end). The change in reading frame on codon (c.) 157 creates a stop codon in new codon 169 affecting protein products initiating translation upstream of c.157, such as FLp53, Δ40p53 or Δ133p53, which become truncated and will lack the HA tag. These proteins and any putative peptides resulting from their cleavage will no longer be detectable using HA antibody. For a better visualization of the shorter p53 forms we treated the cells with Thapsigargin (TH), which activates Δ133p53 and Δ160p53 expression *via* the integrated stress response (ISR)^13^. As expected, under these conditions all alpha isoforms were easily visualized with the HA antibody and R248Q and R273H mutations further increased steady-state levels of FLp53^23^ and shorter isoforms including Δ133p53 and Δ160p53^6^, while the frameshift mutation made the products above Δ160/Δ169p53 undetectable with anti-HA (**Fig. 1D**, left panel). Antibody against FLAG revealed exclusively p53 protein products containing the N-terminus, including the new product with 168 amino acids (1-168) resulting from the frameshift mutation at codon 157 (**Fig. 1D**, right panel). Interestingly, the three top bands under Δ160/Δ169p53 in the HA blots, of approximately 26, 21 and 19 kDa, also almost completely disappeared in the presence of fs-157, though the Δ160/Δ169p53 bands and the band at 18 kDa remained largely unchanged. With this result we can conclude that the three higher bands are C-terminal fragments deriving from the cleavage of larger p53 proteins and that the band migrating around 18 kDa either results exclusively from the processing of Δ160/Δ169p53 isoforms or is directly translated from a TIS downstream of TIS-169.

**Fig. 1.**
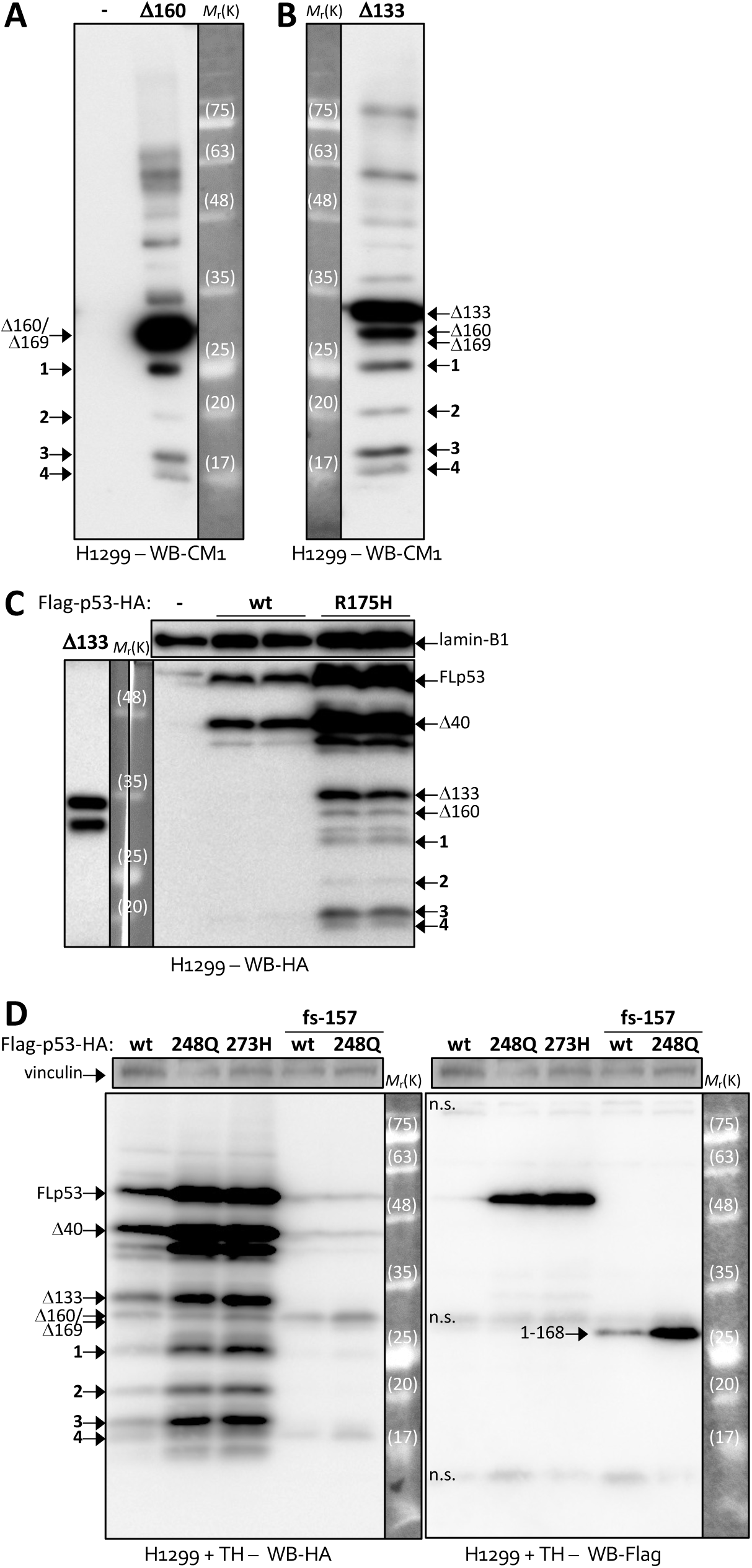
Characterization of short p53 peptides. Western blotting (WB) of p53-negative H1299 cells expressing p53 isoforms Δ160p53 (**A**), Δ133p53 (**B**), full-length (FL) FLAG (N-terminus) and HA (C-terminus) tagged wild-type (wt) or FL tagged mutant R175H (**C**) or R248Q or R273H and with or without a frameshift mutation in codon 157 (fs-157) and treated with integrated stress response inducer drug thapsigargin for 16 h (TH) (**D**) using polyclonal anti-p53 antibody CM1, anti-HA or anti-FLAG antibodies as indicated. Shown are representative data of at least three independent experiments.

### Δ246p53 is translated from a conserved initiation site in codon 246

We investigated the possibility of a new active translation initiation site. Sequence analysis of the 3’-end translated region of human *p53* revealed an *in frame* AUG in codon 246 that is preceded by a strong Kozak sequence (**Fig. 2A**). Prediction software NetStart 1.0 indicates likelihood of translation initiation at that site (values > 0.5) (**Fig. 2B**). Intriguingly, of 50 species analysed, 49 (all vertebrates), from sea lamprey to humans, possess this AUG and a Kozak sequence around it, the only exception was the ascidian sea vase (**Supplementary Fig. S1**). AUG-243, just upstream of c. 246’s Kozak, also showed excellent conservation (96%), suggesting that the c. 243-246 sequence played an important role in *p53* function/regulation throughout evolution. In order to verify if this sequence is at the origin of the 18 kDa band, we mutated both AUG codons (Δ160-243/246A) and expressed, aside, the 246-393 C-terminal amino acid sequence of p53 for size confirmation. 246-393 protein run at exactly the same speed as the 18 kDa band and Δ160p53-M243A/M246A double mutant completely lost the band in question, proving that c.243-c.246 constitutes a TIS and that this encodes for a short C-terminal p53 form of approximately 18 kDa that we termed Δ246p53 (**Fig. 2C**). We have recently shown that a double TIS strengthens the translation of Δ160p53 under different cell conditions^18^, so next we tested if both c.243 and c.246 play an active role in Δ246p53 translation. By mutating each AUG separately or simultaneously in transiently transfected cells, we could see that indeed both seem to contribute for the translation of this “mini” p53 isoform, though it was also clear that TIS-246 has the strongest control over it (**Fig. 2D**), in line with the translation prediction results. We have thus identified a new translation initiation site (TIS) in codon 246 that leads to the expression of a new protein product, Δ246p53, from short p53 transcripts like Δ133p53 mRNA or from activated (through stress stimuli, for example, **Fig. 1D**) or mutated (R175H, R248Q or R273H) full-length p53 mRNA.

**Fig. 2.**
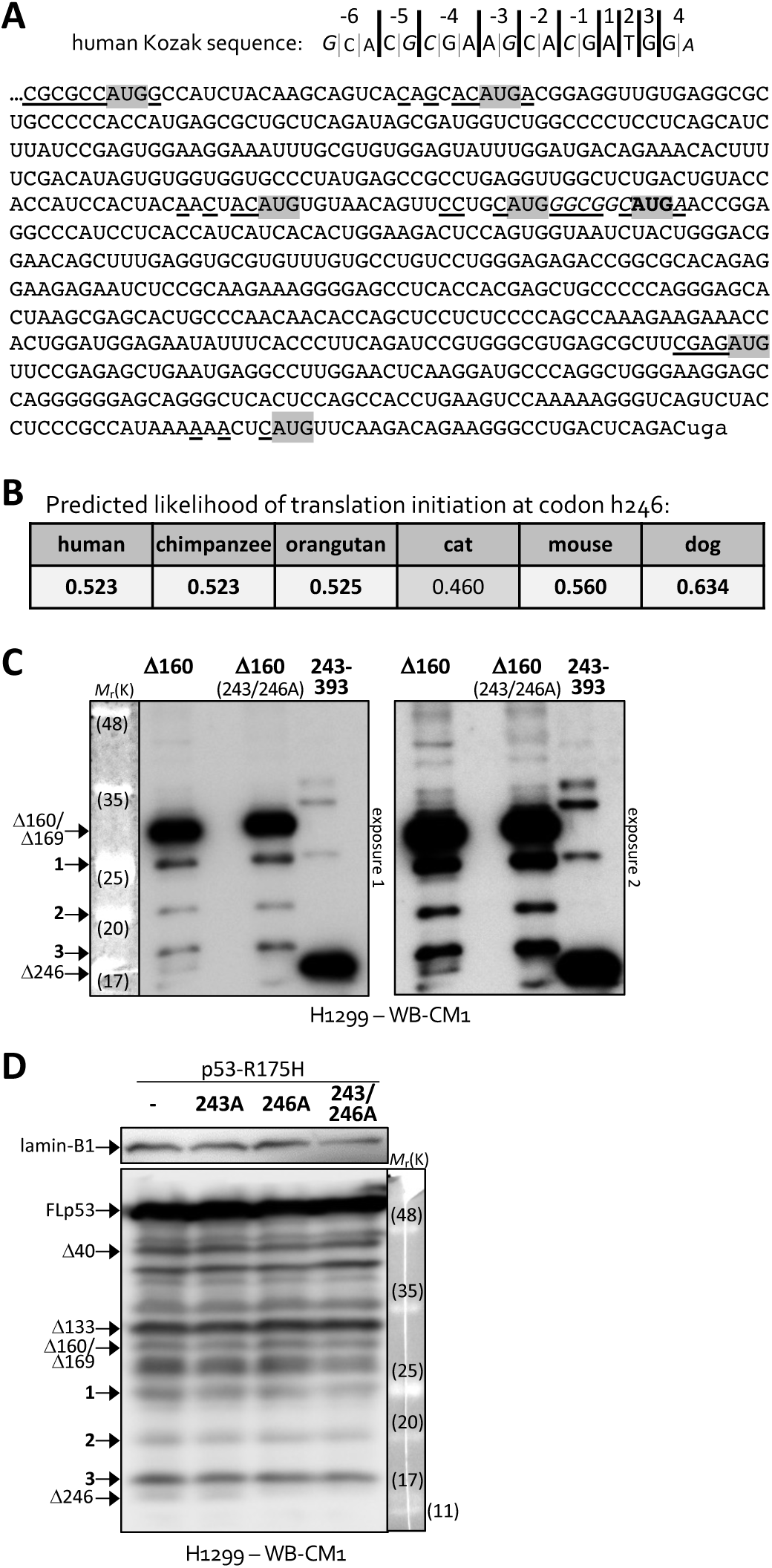
Identification of new p53 isoform, Δ246p53. **A** Most common nucleotides appearing at each position within the Kozak sequence for 10,012 human genes^40^ (top) and 3’ coding region (codons 158-393) of *p53* mRNA (bottom). Lowercase, stop codon; grey boxes, all AUG codons that are *in frame* with the first codon of *p53*; underlined, nucleotides that match the human Kozak sequence shown above; bold, codon 246; italic; Kozak sequence for codon 246. **B** Predicted likelihood of translation initiation from human 246 codon homologs in selected mammal species as calculated by NetStart 1.0. **C, D** Western blotting of p53-negative H1299 cells expressing a C-terminal fragment of p53 (coding sequence 243-393, translating into amino acids 246-393), wt Δ160p53 isoform or Δ160p53 with a double mutation (M243/246A) (**C**) or FL R175H p53 with or without M243A, M246A, or double M243/246A mutation (**D**). CM1, polyclonal anti-p53 antibody. Shown are representative data of at least three independent experiments.

### Δ246p53 is expressed endogenously and induced in response to DNA damage

Δ246p53 was also easily observed in human embryonic kidney 293 (HEK293T) cells (**Fig. 3A**). Its identity was directly confirmed by its size (∼18 kDa), by the same pattern of 4 bands observed in previous blots, and by different antibodies targeting its N-terminus (DO12, aa. 256-267), its C-terminus (Bp53.10, aa. 374-378 or Pab421, aa. 376-378) or, as negative control, an epitope laying just upstream of the Δ246p53 sequence (Pab240, aa. 213-217). Δ246p53 was also detected in a wide variety of cancer cell lines: LN229, HCT116, U2OS, A549, HT1080 and SW480 (**Figs. 3B, C, D** and **Supplementary Fig. S2**); particularly after DNA damage following treatments with Etoposide (ETO), Oxaliplatin (OXA) or Paclitaxel (PTX) (**Figs. 3A-D**). Thapsigargin and Tunicamycin (TUN), which cause endoplasmic reticulum (ER) stress and activate ISR, were not effective in activating Δ246p53 expression in HEK293T or LN229 cells. Δ246p53, and other p53 isoforms including FLp53, were efficiently silenced by different siRNAs targeting p53 mRNA in different cell lines (**Figs. 3E, F** and **Supplementary Fig. S3**). A more specific knock-down was achieved with antisense morpholino oligo (MO) in replacement of siRNA. The MO was designed to bind on top of c.243-c.246 in order to inhibit Δ246p53 translation and though it wasn’t as effective as the siRNA it reduced Δ246p53 levels by 60% with little effect on FLp53 (**Fig. 3F**, right side). Of interest, Δ133p53 and Δ160p53 isoforms were also downregulated by MO. This is because TIS-246 overlaps with the Internal Ribosome Entry Site (IRES) that directs the translation of those two isoforms^13^. Notably, Δ40p53 was also strongly knocked-down by MO, which we think may be related to the fact that Δ40p53 expression is also IRES-dependent^8^.

**Fig. 3.**
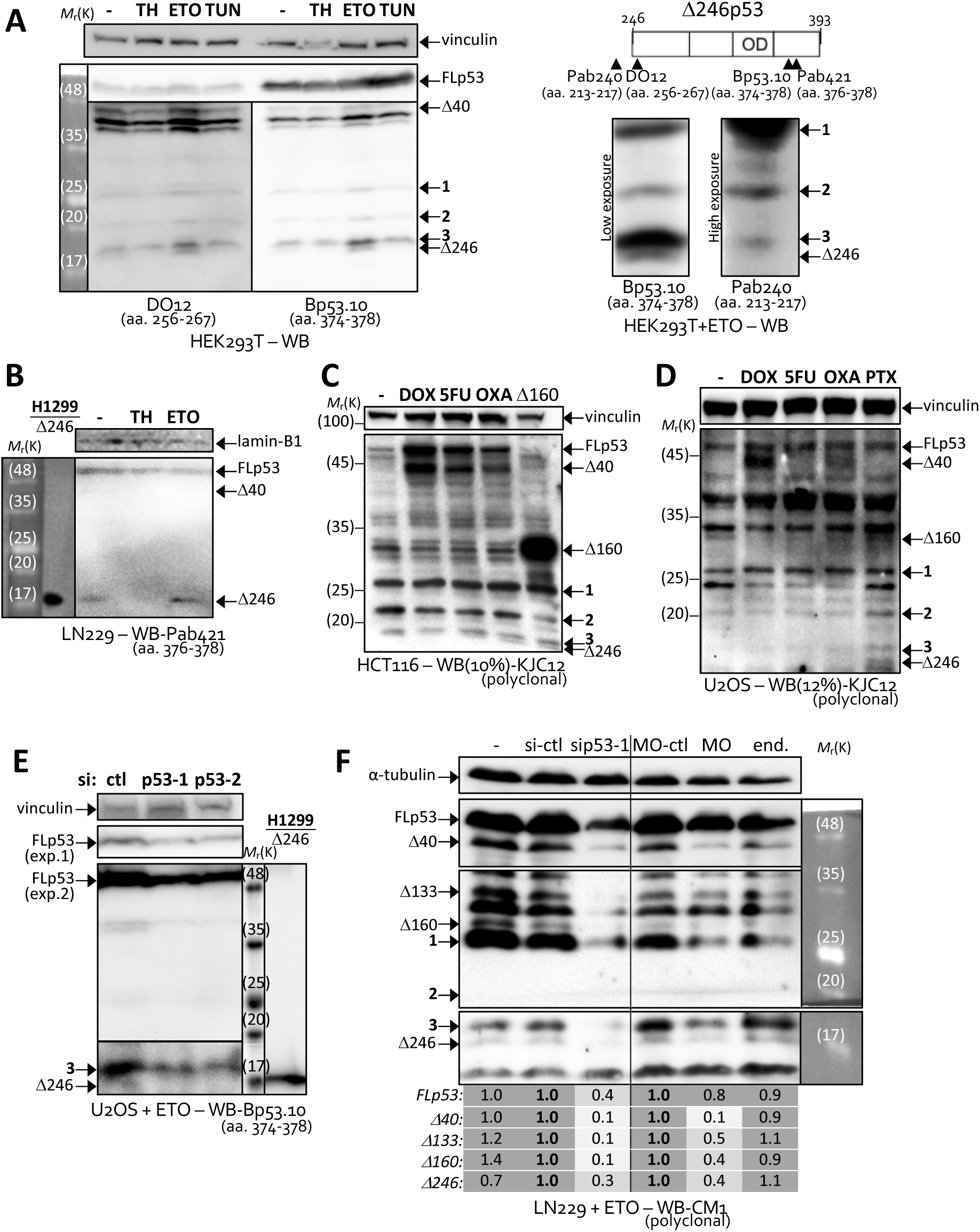
Δ246p53 induction and knock-down in different cell lines. Western blotting (WB) of embryonic kidney HEK293T (**A**), brain glioblastoma LN229 (**B, F**), colorectal carcinoma HCT116 (**C**) and bone osteosarcoma U2OS (**D**) cells treated or not with DNA damaging agents Etoposide (ETO, 5 μM for 21 h except in **B**, 1 μM for 16 h), Doxorubicin (DOX, 0.5 μM for 24 h), 5-Fluorouracil (5FU, 10 μM for 24 h), Oxaliplatin (OXA, 5 μM for 24 h) or Paclitaxel (PTX; 6 μM for 24h) (**A-F**), or integrated stress response inducing drugs Thapsigargin (TH, 16 h) (**A, B**) or Tunicamycin (TUN, 16 h) (**A**) and treated or not with control siRNA (si-ctl) or siRNA against exons 7 or 11 (3’-UTR) of *p53* mRNA (sip53-1 and sip53-2, respectively) or control morpholino oligo (MO-ctl) or MO against TIS-246 (MO) or Endo-Porter alone (end., MO delivery reagent) (**E, F**), as indicated (see also supplementary Figs. S2, S3). DO12, monoclonal anti-p53 (aa 256-267) antibody; Bp53.10, monoclonal anti-p53 (aa 374-378) antibody; Pab240, monoclonal anti-p53 (213-217) antibody; Pab421, monoclonal anti-p53 (aa 376-378) antibody; KJC12, polyclonal anti-p53 (pantropic) antibody; CM1, polyclonal anti-p53 (pantropic) antibody; exp., exposure. Shown are representative data of at least three independent experiments. The numbers under the WB in (**F**) specify, for each line, the amounts of protein for the indicated bands relative to bands showing numbers in bold (either si-ctl, left panel, or MO-ctl, right panel) according to WB quantifications and normalization to α-tubulin.

### Δ246p53 supports senescence and inhibits tumour growth

During our experiments we noticed that on occasion Δ246p53-expressing cells would not divide as much as control cells, so we established cells stably expressing Δ246p53 (**Supplementary Fig. S4A**), plated them, exposed them to DNA-damaging agent Etoposide (ETO) and quantified their numbers daily for three days. We did the same with control cells lacking Δ246p53. Δ246p53 expression was stable throughout the assay (**Supplementary Fig. S4B**) and led to a slight decrease in proliferation/survival, though the difference was not significative (**Supplementary Fig. S4C**). Curiously, the diminution in cell number did not match the total metabolic activity measured by MTT reduction, which failed to indicate any decrease in metabolism for Δ246p53-expressing cells (**Supplementary Fig. S4D**). This could be an indication that cells stopped dividing but remained metabolically active, so we next considered if Δ246p53 could play a role in senescence, as senescent cells may continue enlargement in the absence of cell division^24^. We cultured cells for several days and then weeks and tested them for senescence at different stages using the β-galactosidase assay. While in the first days of culture there was little difference between Δ246p53 and control (**Fig. 4A**), after two weeks, when dishes became overly confluent, Δ246p53 showed a clear inducive effect on the number of senescent cells (**Figs. 4B, C**). Western blotting (WB) analysis confirmed that cells still expressed Δ246p53 at high density (**Fig. 4D**). Since epithelial cells, such as H1299, more preferentially respond to stress *via* cell death, while fibroblasts more often activate senescence pathways under those conditions^25^, we next tested the effect of Δ246p53 on the activation of senescence using fibroblast-like mesenchyme-derived fibrosarcoma cells HT1080. Indeed, genotoxic stress using Oxaliplatin effectively activated *p21* as well as Δ246p53 expression in HT1080 cells (**Fig. 4E**) and β-galactosidase assay showed a 6-fold increase in senescence in the presence of Δ246p53 (**Fig. 4F**). Δ246p53’s role in cell-cycle control was also confirmed by Fluorescence-Activated Cell Sorting (FACS) (**Fig. 4G**), which revealed an increase in the number of cells in G_0_/G_1_, the phase most associated with senescence (**Figs. 4G, H**).

**Fig. 4.**
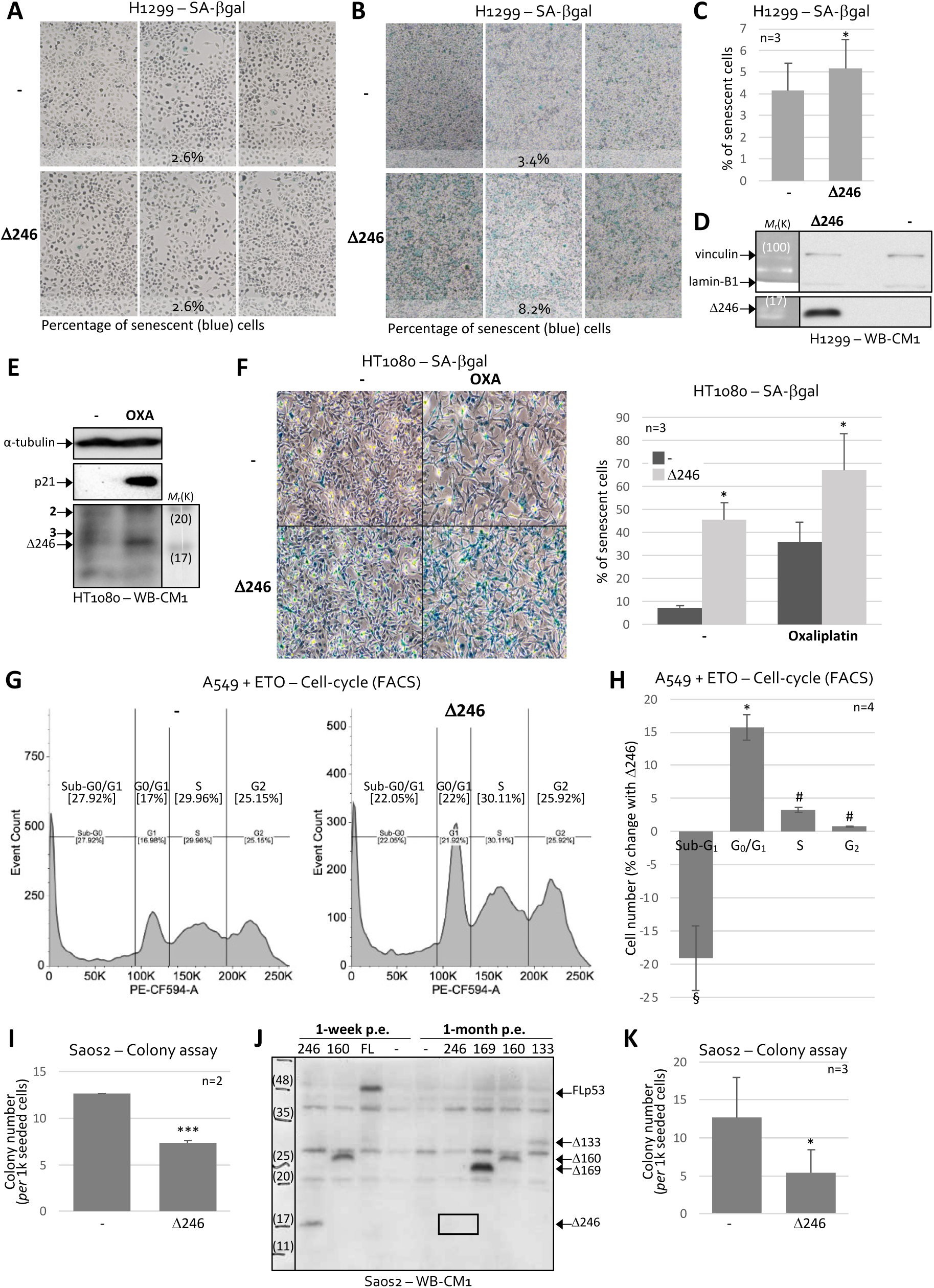
Δ246p53’s functions in senescence and tumour suppression. **A-C** Senescent H1299 cells (blue) expressing or not Δ246p53 and cultured under- (**A**) or over- (**B**) confluent, detected by X-Gal cleavage. Percentage of blue staining is indicated at the bottom of each panel of three photographs and averages of 81 photographs (3 independent experiments) are shown in (**C**). **D** WB of H1299 cells stably expressing or not Δ246p53 after over-confluent culture shown in (**B**). **E** WB of HT1080 fibrosarcoma cells treated or not with Oxaliplatin (OXA). **F** Senescent HT1080 cells (blue) expressing or not Δ246p53 and treated or not with Oxaliplatin, detected by X-Gal cleavage. Percentage values of blue staining are indicated as averages of 81 photographs (right panel). **G, H** Fluorescence-activated cell sorting (FACS)-mediated cell-cycle analyses of A549 cells expressing or not Δ246p53 and treated with etoposide (10 μM for 21 h) and stained with propidium iodide, followed by FlowJo (**G**) and statistical **(H)** analyses. **I, K** Soft agar colony formation assays with *p53*-null Saos2 cells stably expressing or not Δ246p53. **J** WB of Saos2 cells stably expressing or not Δ246p53 and other p53 forms (as indicated) 1-week or 1-month post-establishment (p.e.). Shown are averages ± s.d. of n experiments as indicated or representative data of at least three independent experiments (#P > 0.1, §P < 0.1, *P < 0.05 and ***P < 0.005 compared to cells not expressing Δ246p53). CM1, polyclonal anti-p53 antibody.

Next, the density-dependent activity of Δ246p53 observed in the senescence assays prompted us to investigate its ability to affect colony formation in soft agar. The capacity for anchorage-independent growth in soft agar is considered a hallmark of carcinogenesis^26^, and the competence to regulate it is often used as a defining feature of oncogenes (if promoting) or tumour suppressor genes (if repressing). Excitingly, in the first two assays, Δ246p53 reduced the number of colonies to less than 60% of those of the control (**Fig. 4I**). The third assay showed no differences (**Supplementary Fig. S5**), which we then confirmed was because, one-month post-establishment (p.e.), cells that were still proliferating had completely lost Δ246p53 expression (**Fig. 4J**), supporting Δ246p53’s effectiveness in restricting cell division. After establishing new cells (**Supplementary Fig. S6**), Δ246p53’s strong inhibition of colony formation was again observed (**Fig. 4K**). These results suggest that this gene product can act as a tumour suppressor.

### Δ246p53 regulates *HDM2* and *p21* through its interactions with different p53 family proteins

We next wondered how Δ246p53 affects cell condition and colony formation. Several C-terminal isoforms of *p53* have previously been reported to bind to FLp53 and work as regulators, co-factors or activators by lowering/improving/diversifying the transcriptional activation of target genes^13,27^. Δ246p53 preserves the oligomerization domain (OD) present in these isoforms and could in theory also interact with FLp53 and affect its capacity to transactivate target genes (**Fig. 5A**). Indeed, we could successfully co-immunoprecipitate FLp53 by pulling down Δ246p53-HA through an HA tag (**Fig. 5B**). Pulling down FLp53 through an N-terminal epitope (aa 11-25) using DO-1 antibody also efficiently captured Δ246p53, which lacks the epitope, unless it was mutated at aa 342 (R342P, monomer, mono or Δ246p53m), an essential amino acid for the tetramerization and dimerization with other p53 proteins^28^, confirming that the two proteins interact (**Fig. 5C**).

**Fig. 5.**
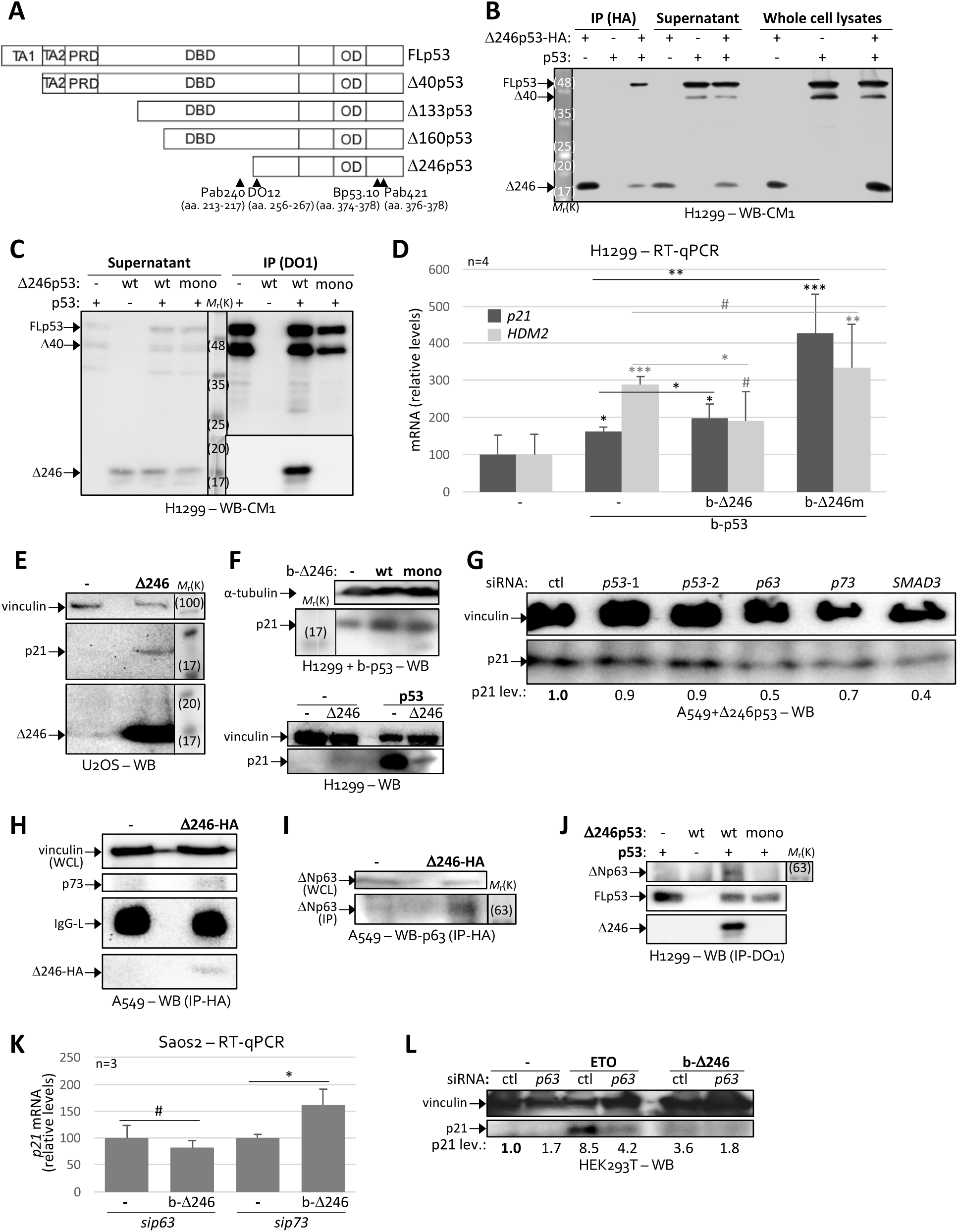
Δ246p53 binding to p53 family members and effect on downstream targets. **A** Representation of alternative translation products of *p53* and their domains. TA, transactivation domain; PRD, proline-rich domain; DBD, DNA-binding domain; OD, oligomerization domain. **B**, **C** Immunoprecipitation (IP) using HA(**B**) or DO1 (**C**) antibody in lysates from H1299 cells expressing or not oligomeric (wt) or monomeric (R342P, mono) Δ246p53, full-length (FL) p53 (p53) or tagged Δ246p53-HA or a combination of these followed by Western blotting (WB) as indicated. Whole cell lysates (WCL) shown in WB were recovered before IP. Supernatant of samples was recovered after IP and before washes. **D** RT-qPCR of total RNA extracted from H1299 cells expressing or not bicistronic constructs for endogenous-level expression of FLp53 (b-p53), oligomeric Δ246p53 (b-Δ246) or monomeric Δ246p53 (b-Δ246m) as indicated using primers against *p21* or *Hdm2* mRNAs and normalized against *GAPDH* and *RPS18* housekeepers’ mRNA levels. **E-G** WB of U2OS (**E**), p53-null H1299 (**F**) or A549 (**G**) cells expressing or not high levels (Δ246) or physiological levels (b-Δ246) of Δ246p53 in its oligomeric (wt) or monomeric (mono) form or high levels (p53) or physiological levels (b-p53) of FLp53 and treated or not with control siRNA (si-ctl) or siRNA against exons 7 or 11 (3’-UTR) of *p53* mRNA (sip53-1 and sip53-2, respectively) or siRNA against *p63*, *p73* or *SMAD3* mRNAs, as indicated. **H-J** IP using HA (**H, I**) or DO1 (**J**) antibody in lysates from A549 (**H, I**) or H1299 (**J**) cells expressing or not oligomeric (wt) or monomeric (R342P, mono) Δ246p53, FLp53 (p53) or tagged Δ246p53-HA or a combination of these followed by Western blotting (WB) as indicated. Whole cell lysates (WCL) shown in WB were recovered before IP. **K** RT-qPCR of total RNA extracted from p53-null Saos2 cells expressing or not bicistronic constructs for endogenous-level expression of Δ246p53 (b-Δ246) and treated with siRNA against *p63* or *p73* mRNAs as indicated using primers against *p21* mRNA and normalized against *GAPDH* and *RPS18* housekeepers’ mRNA levels. **L** WB of HEK293T cells expressing or not endogenous-like levels of Δ246p53 (b-Δ246) and treated or not with genotoxic agent Etoposide (ETO) and with control siRNA (si-ctl) or siRNA against *p63*, as indicated. The numbers under the WB (G, L) specify the amounts of p21 protein for the indicated lanes relative to bands showing numbers in bold (either si-ctl, G, or si-ctl with empty vector (-) samples, L) according to WB quantifications and normalization to vinculin. Shown are averages ± s.d. of n experiments as indicated or representative data of at least three independent experiments (#P > 0.1, *P < 0.05, **P < 0.01 and ***P < 0.005 compared to cells not expressing any p53 or as indicated).

We then verified if Δ246p53 expression affected the transcription of p53 target genes *p21* and *HDM2*, main regulators of p53-mediated senescence and p53 protein stability, respectively^29,30^. Both genes were activated by p53, as expected, even when expressing p53 at very low levels using a bicistronic construct^31^ (b-p53, see **Supplementary Fig. S7A**). However, these genes responded differently to Δ246p53 (also expressed from a bicistronic construct, b-Δ246, to match endogenous levels – represented in **Supplementary Fig. S7A**, levels shown in **Supplementary Fig. S7B**): while *p21* was further enhanced by Δ246p53, *HDM2* was inhibited (**Fig. 5D**). When testing the monomer mutant of Δ246p53 (b-Δ246m), the inhibitory effect on *HDM2* was lost, suggesting that Δ246p53 inhibits *HDM2* by interacting with FLp53 and abrogating its transcription factor function. Curiously, monomeric Δ246p53 was even more efficient in activating *p21* than the WT oligomeric form (**Fig. 5D**). WT Δ246p53 and monomeric Δ246p53 also activated *p21* at the protein level in different cell lines in the presence of low levels of FLp53 (**Figs. 5E** and **5F, top panel**) or even in the absence of FLp53 (**Fig. 5F, bottom panel**, left two lanes), showing that Δ246-mediated induction of *p21* is independent of FLp53. Interestingly, similarly to what was observed for *HDM2*, Δ246p53 strongly inhibited *p21* expression when this one was activated by FLp53 (**Fig. 5F, bottom panel**, right two lanes), which suggests that Δ246p53 is a dominant-negative regulator of FLp53, while activating *p21* through a different path.

We have previously observed that many – if not all – of the functions observed for gain-of-function mutants of *p53* are natural functions of different p53 products that have evolved within the *p53 locus* for millions of years throughout the development of major animal *phyla* like mammals (Δ160p53), primates (Δ133p53) and possibly vertebrates (Δ246p53)^6,18,32^. Rather than creating new functions that did not previously exist (neomorphic), these mutations seem to be activating existing functions – and thus the gain-of-function effect – that are specific of the less studied p53 products. With that in mind, and in an effort to identify the factor(s) involved in Δ246p53-mediated senescence, we assembled a table of potential interacting-partners of Δ246p53 based on their known involvement in ageing and *p21* activation as well as interaction with some form of mutant p53 (**Table 1**). We next used AlphaFold2 to help us estimate which of these factors is more likely to interact with Δ246p53, even if failing to bind to wild-type FLp53 (**Supplementary Fig. S8**). The best hits were clearly p73 and p63, followed by Smad3 (marked with green borders). So, we tested if any of these three genes had an impact on p21 levels in the presence of Δ246p53, by targeting them with siRNA (**Fig. 5G**). We also targeted *p53* and confirmed it had little effect on p21 levels in the presence of Δ246p53. All the others, however, were necessary to maintain high levels of p21 protein (**Fig. 5G**). We next pulled-down Δ246p53 and investigated if it bound to any of these factors. While the interaction with p73 was not clear (**Fig. 5H**), we observed co-immunoprecipitation of ΔNp63 (**Fig. 5I**). Remarkably, FLp53 also captured ΔNp63 when immunoprecipitated with DO-1 but only in the presence of Δ246p53 (**Fig. 5J**). Abolishing FLp53-Δ246p53 interaction using the monomeric mutation R342P (Δ246p53-mono) led to a complete loss of binding to ΔNp63, showing that it is Δ246p53 that mediates the contact between FLp53 and ΔNp63.

**Table 1.**
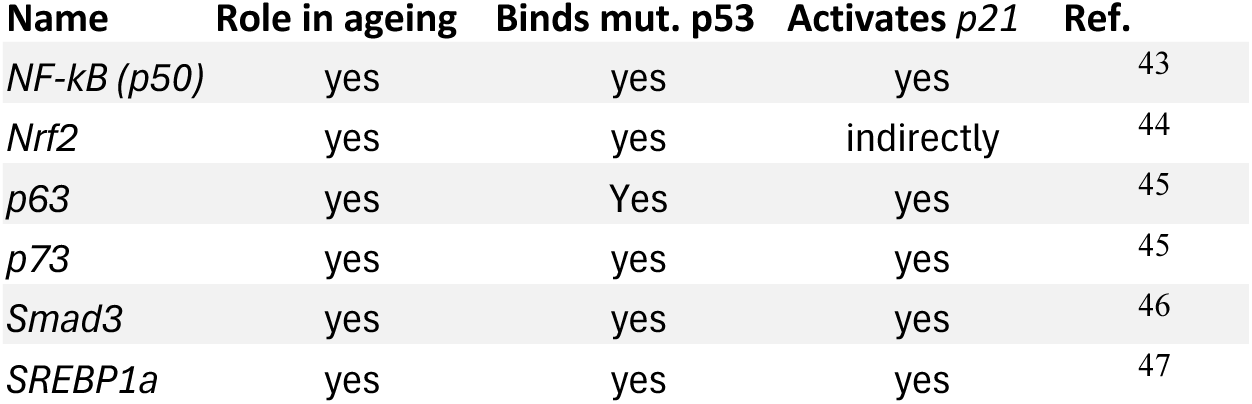
Potential Δ246p53–binding partners that bind mutant p53, activate *p21* and have a known impact on ageing.

| **Name** | **Role in ageing** | **Binds mut. p53** | **Activates *p21*** | **Ref.** |
| --- | --- | --- | --- | --- |
| *NF-kB (p50)* | yes | yes | yes | 43 |
| *Nrf2* | yes | yes | indirectly | 44 |
| *p63* | yes | Yes | yes | 45 |
| *p73* | yes | yes | yes | 45 |
| *Smad3* | yes | yes | yes | 46 |
| *SREBP1a* | yes | yes | yes | 47 |

To test if ΔNp63 is involved in the effect that Δ246p53 exerts on *p21* expression, we knocked-down p63 using siRNA (**Supplementary Fig. S9**) and then quantified the levels of *p21* mRNA (**Fig. 5K**) and p21 protein (**Fig. 5L**) in the presence or absence of physiological Δ246p53 levels (b-Δ246). While siRNAs against *p73* (**Fig. 5K**) or *SMAD3* (**Supplementary Fig. S10**) did not prevent Δ246p53-mediated induction of *p21*, knock-down of *p63* blocked the isoform’s action (**Fig. 5K, L, Supplementary Fig. S11**), suggesting that *p63* is a major player in the observed phenotype. Altogether, our data indicate that, mechanistically, Δ246p53 forms complexes with FLp53 and ΔNp63 leading to a dominant-negative effect on FLp53’s activation of downstream targets *HDM2* and *p21*, while simultaneously promoting p63-mediated activation of *p21* (a model of action is proposed in **Fig. 6** and discussed in the following section).

**Fig. 6.**
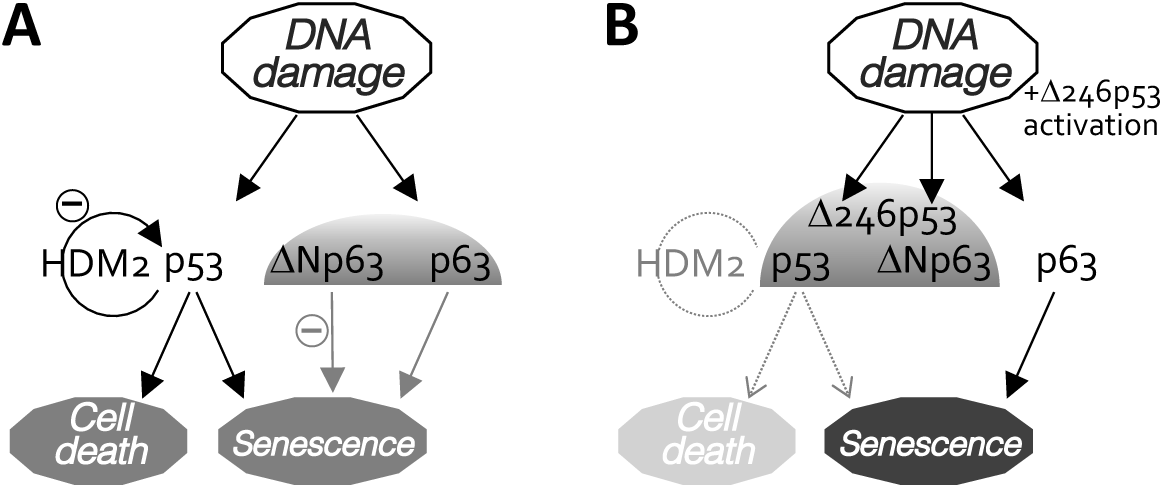
Proposed model of Δ246p53 action following induction by DNA damage. **A** In the absence of Δ246p53, and in response to genotoxic stress: FLp53 (p53) activates senescence through *p21* but also cell death, as well as a negative feedback-loop (-) with HDM2; TAp63 (p63) activates senescence, though its action may be partially inhibited by dominant-negative ΔNp63 isoform; ΔNp63 inhibits *p21* directly at its promoter. **B** In the presence of Δ246p53: FLp53 (p53) and ΔNp63 are both bound by Δ246p53, which restricts their functions, including in the activation of cell death and senescence; the *p63-p21* branch that regulates senescence becomes strongly activated.

## DISCUSSION

Here we identified a new naturally occurring protein isoform of *p53*, Δ246p53. Δ246p53 was detected in cell lines of embryonic kidney origin (HEK293T), brain glioblastoma (LN229), colorectal carcinoma (HCT116), bone osteosarcoma (U2OS) (**Fig. 3**), lung adenocarcinoma (A549), fibrosarcoma (HT1080), and colon adenocarcinoma (SW480) (**Supplementary Fig. S2**). These include non-cancer (HEK293T) and cancer cells with wild-type (HEK293T, U2OS, A549, HT1080 and HCT116) or mutant (LN229 [P98L] and SW480 [R273H/P309S]) *p53*. We failed to detect Δ246p53 in breast biopsies from cancer patients using WB (n=14, data not shown). We think this may be related to its tumour suppressor function. Δ246p53 expression was enhanced following DNA damage, when it reached levels sometimes comparable or superior to Δ40p53, the first described and well-established shorter human p53 isoform^33^ (**Figs. 3A-D**).

Δ246p53 is expressed from a translation initiation site (TIS) in codon 246 and lacks both transactivation domains (TA), the proline rich domain (PRD) and most of the DNA binding domain (DBD) (**Fig. 5A**). It retains the oligomerization domain (OD) through which p53 proteins homo-oligomerize in order to transactivate target genes^34^. We could confirm Δ246p53’s interaction with full-length (FL) p53 protein (**Fig. 5B, C**) as well as the role of the OD in this interaction using a monomerizing mutation within this domain (R342P)^28^ (**Fig. 5C**). Functionally, this bond led to a dominant-negative effect on the transactivation of FLp53 target genes *HDM2* (**Fig. 5D**) and *p21* (**Fig. 5F, bottom panel**). Interestingly, in conditions when FLp53 was absent or inactive (p53-null cells or non-stressed cells), Δ246p53 activated *p21* independently of FLp53 (**Figs. 5D-F**). A small *in silico*-siRNA–combined screening of *p21* activators known to interact with mutant p53 followed by co-immunoprecipitation assays led to the identification of *p63* as a key mediator of Δ246p53’s function (**Figs. 5G-L**, **Table 1**, **Supplementary Figs. S8-11**). The interaction we observed was not between Δ246p53 and TAp63 but between Δ246p53 and ΔNp63, which could again explain the outcome on p21 levels through a dominant-negative effect. While TAp63 has an important p53-independent role in senescence and ageing that relies on *p21* activation^35,36^, ΔNp63 binds directly to the *p21* promoter to inhibit its activation^37^ and has a reported negative-dominant effect on TAp63^38^. So, we propose that Δ246p53 forms a complex with FLp53 and ΔNp63 (observed in **Figs. B, C, I, J**) and inactivates both, hampering FLp53-dependent activation of *HDM2* and *p21* and likely also preventing ΔNp63-mediated inhibition of *p21* and TAp63, which will in turn trigger senescence (see proposed model in **Fig. 6**). In fact, *p63* was required for Δ246p53-mediated induction of *p21* (**Figs. 5K, L**). Of note, these experiments (**Figs. 5 D, F, K, L**) were performed with bicistronic constructs that express very low levels of Δ246p53 through internal translation initiation^8^, similar to endogenous expression (**Supplementary Fig. S7**, b-Δ**246**), closely mimicking the natural conditions in the cell. The effect observed on *p21* is consistent with increased restrictions on cell-cycle progression. In fact, Δ246p53 induced G_0_/G_1_ arrest (**Figs. 4G, H**), senescence (**Figs. 4A-F**) and impaired tumour colony formation in soft agar (**Fig. 4I-K**). The role of Δ246p53 on senescence was more clearly observed in fibroblast-like mesenchyme-derived fibrosarcoma cells HT1080 (**Fig. 4F**), than in lung carcinoma H1299 cells of epithelial origin (**Fig. 4A-C**). This may be related to the fact that epithelial cells more preferentially respond to stress *via* cell death, while fibroblasts more often activate senescence pathways under those conditions^25^. The effect of Δ246p53 on total cell number was not so evident (**Supplementary Fig. S4C**), and this may be because the surge in senescence seems to be accompanied by a reduction in cell death (Sub-G_0_/G_1_ populations in **Fig. 4G, H**; observe also the effect of Δ246p53 on cell number in the presence of Oxaliplatin in **Fig. 4F, left panel**; also included in our model in **Fig. 6**).

Throughout our characterization of Δ246p53, we have also identified 3 common C-terminal cleavage products of p53, of about 26 (#1), 21 (#2) and 19 (#3) kDa. Though we did not identify the exact cleavage sites, we noticed that Δ160/Δ169p53 isoform produces predominantly #1, while Δ133p53 and FLp53 create larger amounts of #3 (compare the intensity of the two fragments in **Fig. 1A-C**). In fact, when the contributions from Δ133p53 and FLp53 were masked by the frameshift mutation in codon 157, #1 became the main cleavage product (**Fig. 2D**), as observed with Δ160p53 expression (**Fig. 1A**). This fragment is possibly the result of caspase-mediated cleavage at amino acid (aa) 186^4^. Cleavage at this site creates an N-terminal fragment of about 36 kDa^4^ also observed in our blots when using N-terminal or polyclonal antibodies (**Fig. 2D** and not shown). These fragments were shown to induce mitochondrial membrane depolarization^4^, whereas the roles of peptides #2 and #3 remain to be investigated.

Noticeably, the natural protein that we have identified here as a product of TIS-246 has been “created” and studied previously, under the name of M protein^39^ – M for mutant in this case, not mini. Importantly, Moore *et al*. made some similar observations to our findings: M protein binds to FLp53 but also displayed FLp53-independent growth suppression, which we likewise documented for Δ246p53 (**Figs. 5C** and **4K**). Donehower’s group also reported a stabilizing and activating effect on FLp53. We did not observe these, but we think it could be expected under some conditions due to the inhibition of the negative regulator E3-ubiquitin ligase Hdm2. In a previous publication the same group presented the mutant p53 mice, *p53^+/m^*, where ^+^ stands for wild-type (WT) and *^m^* stands for a mutant allele in which most of the *p53* gene is deleted and translation (tested *in vitro* in this study) initiates from mouse codon 243, which corresponds to human codon 246 (so the start codon for M protein/Δ246p53) (see **Supplementary Fig. S1** for sequence comparison)^15^. Even though a larger part of the chromosome was deleted in this *m* allele – not just *p53* – the authors attributed most of the phenotype to the expression of the M protein (Δ246p53). Effectively, as we saw here for Δ246p53 (**Figs. 4 and 5**), the *m* allele activated *p21* expression and senescence. More spectacularly, *p53^+/m^*mice exhibited resistance to tumour formation – that we also observe *in cellula* for Δ246p53 (**Figs. 4I-K**) – and aged prematurely. Future research will be needed to more fully define the contribution of Δ246p53 to tumour suppression and ageing.

## ACKNOWLEDGMENTS

This work was supported mainly by grants Ishizue-2021 and Ishizue-2023 (Kyoto University) and the European Union’s Horizon 2020 research and innovation programme under grant agreement No 857524, as well as grants PTDC/MED-ONC/32048/2017 and PTDC/BIM-ONC/4890/2014 from the Portuguese Foundation for Science and Technology (Fundação para a Ciência e a Tecnologia, FCT), grant 18K07229 (KAKENHI) from Japan Society for the Promotion of Science (JSPS), a grant from Takeda Science Foundation (2017 Medical Research Fellowship (Oncology, Basic)) and a grant from Astellas Research Foundation for Pathophysiology and Metabolism (project number 203180600044) attributed to M.M.C; and also UID/00100 BioISI (DOI: 10.54499/UIDB/04046/2020) centre grant from FCT (Portugal) and funds from Instituto Nacional de Saúde Doutor Ricardo Jorge. S.N.P. and M.J.L.I. were partially supported by Otsuka Toshimi Scholarship Foundation. S.N.P. was also supported by Monbukagakusho Honors Scholarship and 2019 International Students (Learning Encouragement Scholarship). A.C.R. was the recipient of a PhD scholarship from the Portuguese Foundation for Science and Technology (2020.06982.BD). p53 antibodies Pab421, Bp53.10, CM1 and DO12 were a gift from Bořivoj Vojtěšek (Masaryk Memorial Cancer Institute, Brno, Czech Republic). p53 antibody KJC12 was a gift from Jean-Christophe Bourdon (University of Dundee, Dundee, United Kingdom) to Y.C.. We also thank Makoto Hayashi (Kyoto University) for providing usage access to StepOnePlus (Applied Biosystems) and other equipment (light and fluorescent microscopy) and reagents, and Shohab Youssefian, Minsoo Kim and Dean Thumkeo (Kyoto University) for reagents as well.

## AUTHOR CONTRIBUTIONS

SNP, ACR and MJLI conducted and supervised experiments. JZ, KK, RS, VD and FSR also conducted experiments. YC, AB and LR provided essential reagents. MMC conceptualized and supervised the study and all experiments. MMC and SNP wrote the paper. All authors were involved in analyses, interpretation and discussion of data. All authors reviewed and were given a chance to comment on the paper.

---

**Supplementary Fig. S1.**
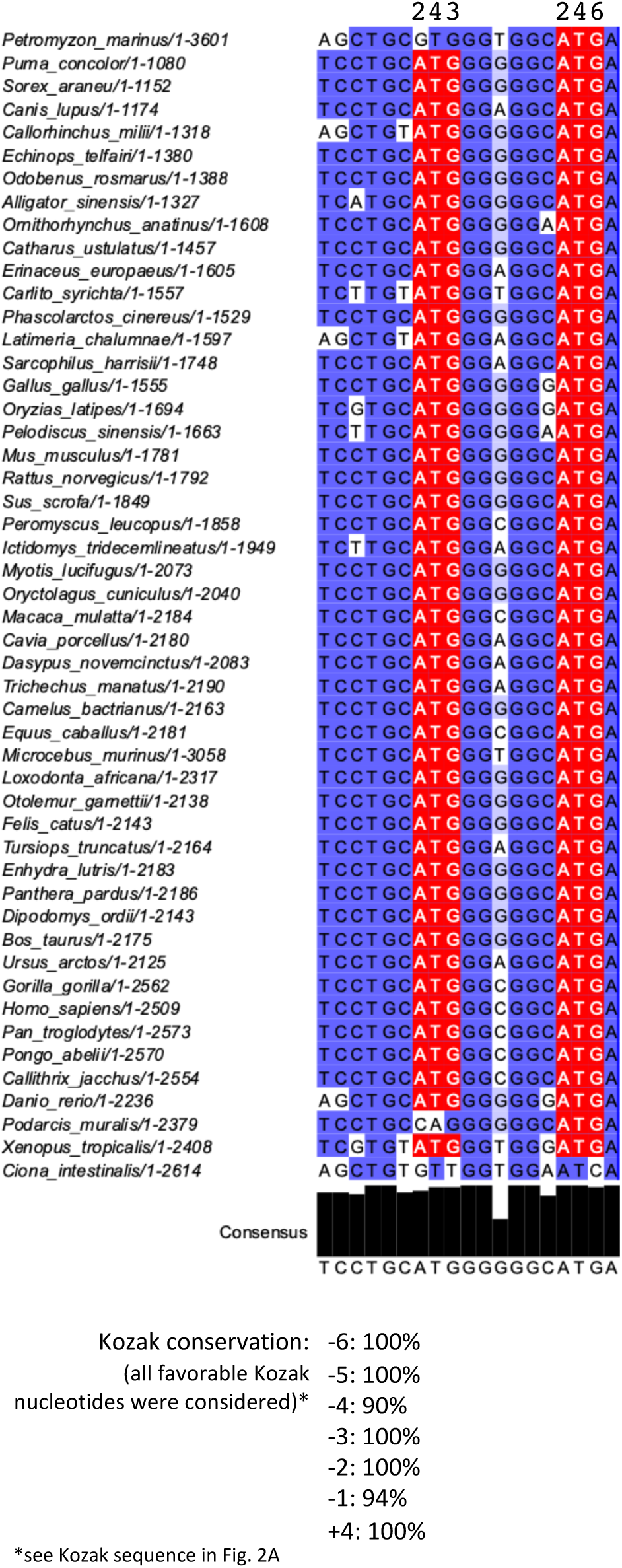
hTIS-246 and its Kozak sequence in different vertebrate species. Sequence alignment of *p53* regions surrounding corresponding human ATG codons 243 and 246 from 50 different species of chordate using MAFFT server within Jalview (top). ATG codons are shown in red and conserved nucleotides in purple. Percentage conservation among the vertebrate species above for each position of the Kozak sequence (bottom).

**Supplementary Fig. S2.**
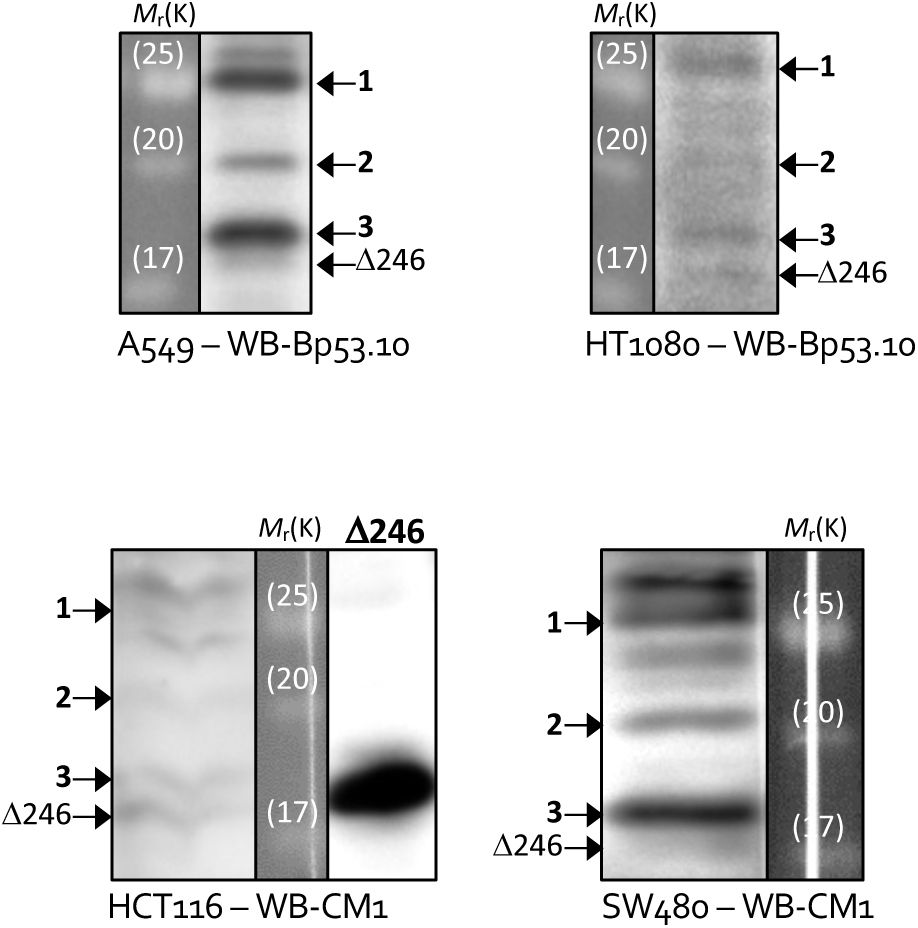
Δ246p53 in different cell lines. Western blotting against p53 in lung adenocarcinoma (A549), fibrosarcoma (HT1080), colorectal carcinoma (HCT116) and colon adenocarcinoma (SW480) cells. Bp53.10, monoclonal anti-p53 (aa 374-378) antibody; CM1, polyclonal anti-p53 antibody. Shown are representative data of at least three independent experiments.

**Supplementary Fig. S3.**
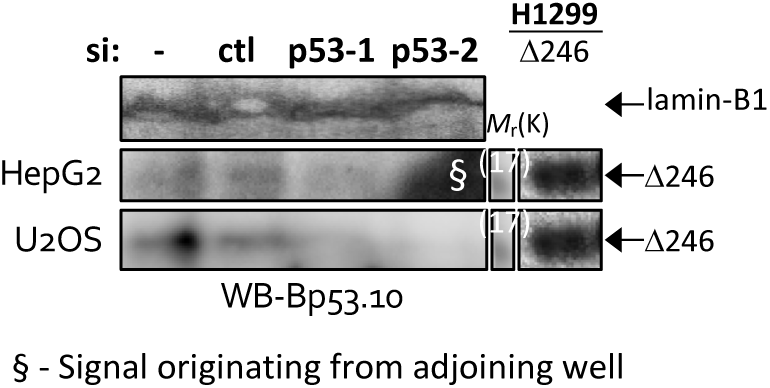
Knock-down of Δ246p53 in different cell lines using two different siRNA. Western blotting (WB) of hepatocellular carcinoma HepG2 and bone osteosarcoma U2OS cells treated or not with control siRNA (si-ctl) or siRNA against exons 7 or 11 (3’-UTR) of *p53* mRNA (sip53-1 and sip53-2, respectively), as indicated. Bp53.10, monoclonal anti-p53 (aa 374-378) antibody. Shown is representative data of three independent experiments.

**Supplementary Fig. S4.**
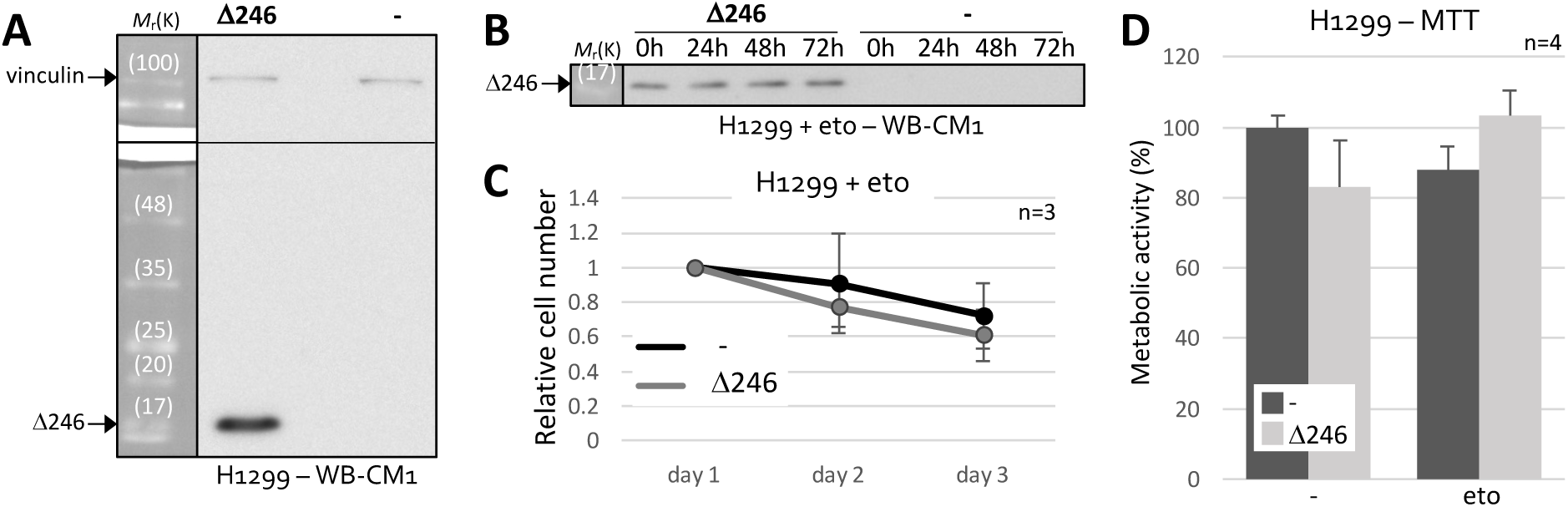
Δ246p53 in proliferation and metabolism. **A, B** Western blotting (WB) of *p53*-null H1299 cells stably expressing or not Δ246p53 (**A, B**) and treated or not with DNA damaging agent etoposite, as indicated (eto) (**B**). **C** Relative cell number following etoposide treatment in H1299 stably expressing or not Δ246p53. **D** MTT assay (metabolic activity) of H1299 cells expressing or not Δ246p53 and treated or not with etoposide (eto, 21 h). Shown are averages ± s.d. of n experiments as indicated or representative data of at least three independent experiments. CM1, polyclonal anti-p53 antibody.

**Supplementary Fig. S5.**
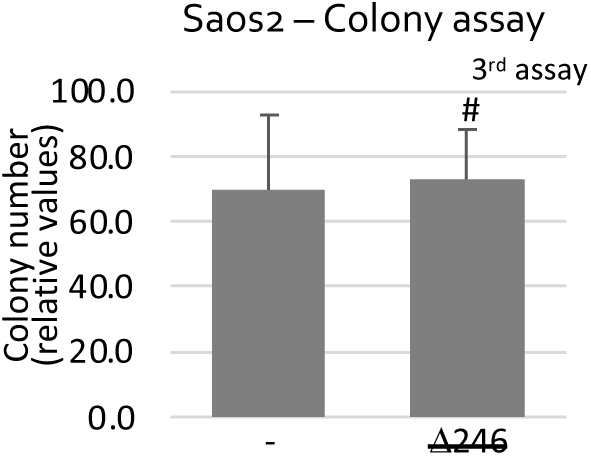
Saos2 cells that lost Δ246p53 expression 1-month post-establishment () no longer showed an effect in soft-agar colony formation assay. Compare with Figs. 4K-M. Shown are averages ± s.d. from 6 replicates, (#P > 0.1).

**Supplementary Fig. S6.**
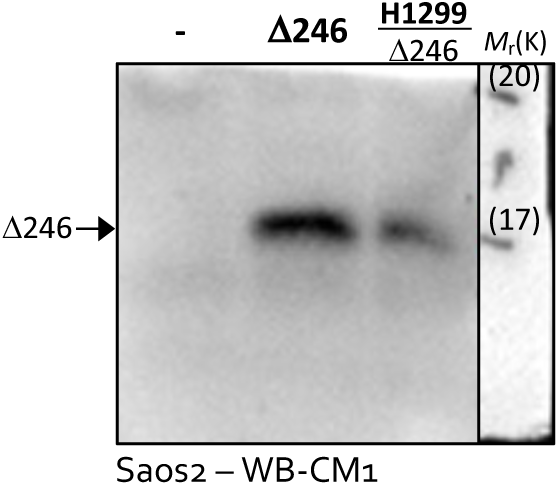
Stable cell lines. Western blotting (WB) of *p53*-null Saos2 cells stably expressing or not Δ246p53. CM1, polyclonal anti-p53 antibody.

**Supplementary Fig. S7.**
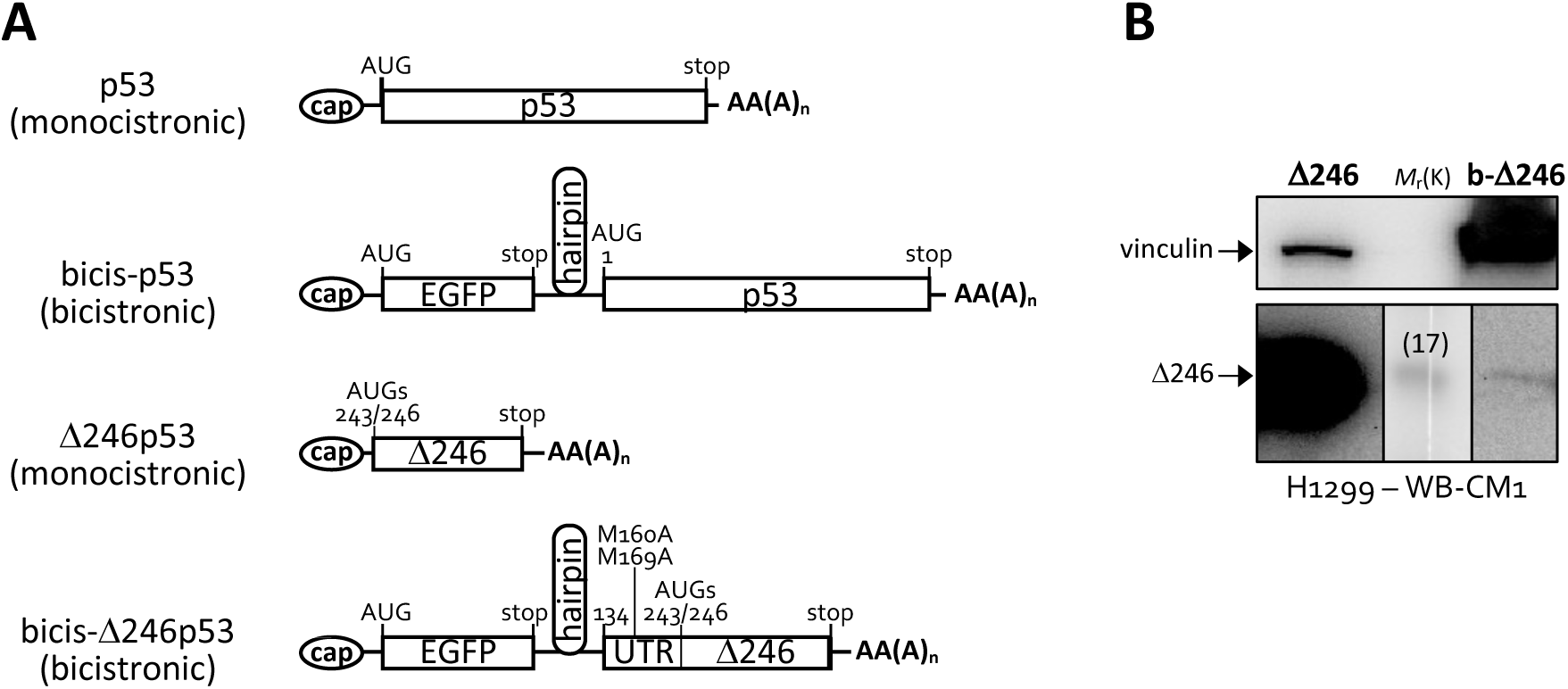
Endogenous-level expression of Δ246p53 and full-length (FL)p53 using bicistronic constructs. **A** Representation of monocistronic and bicistronic mRNA transcripts originating from plasmid constructs used in this study for the expression of high levels of FLp53 (p53) or Δ246p53 (Δ246p53) or low endogenous-like levels of FLp53 (bicis-p53) or Δ246p53 (bicis- Δ246p53. In construct bicis-Δ246p53 the region upstream of the Translation Initiation Sites (TISs) AUG_243_ and AUG_246_ for Δ246p53 (labelled as UTR) was kept because it contains the Internal Ribosome Entry Site (IRES) for Δ246p53 expression; TISs AUG_160_ and AUG_169_ were mutated to prevent translation of Δ160p53/Δ169p53 isoform^41,42^. **B** Western blotting against p53 in *p53*-null H1299 cells expressing monocistronic (Δ246) or bicistronic^8^ (b-Δ246) constructs of Δ246p53. CM1, polyclonal anti-p53 antibody. Shown is representative data of three independent experiments.

**Supplementary Fig. S8.**
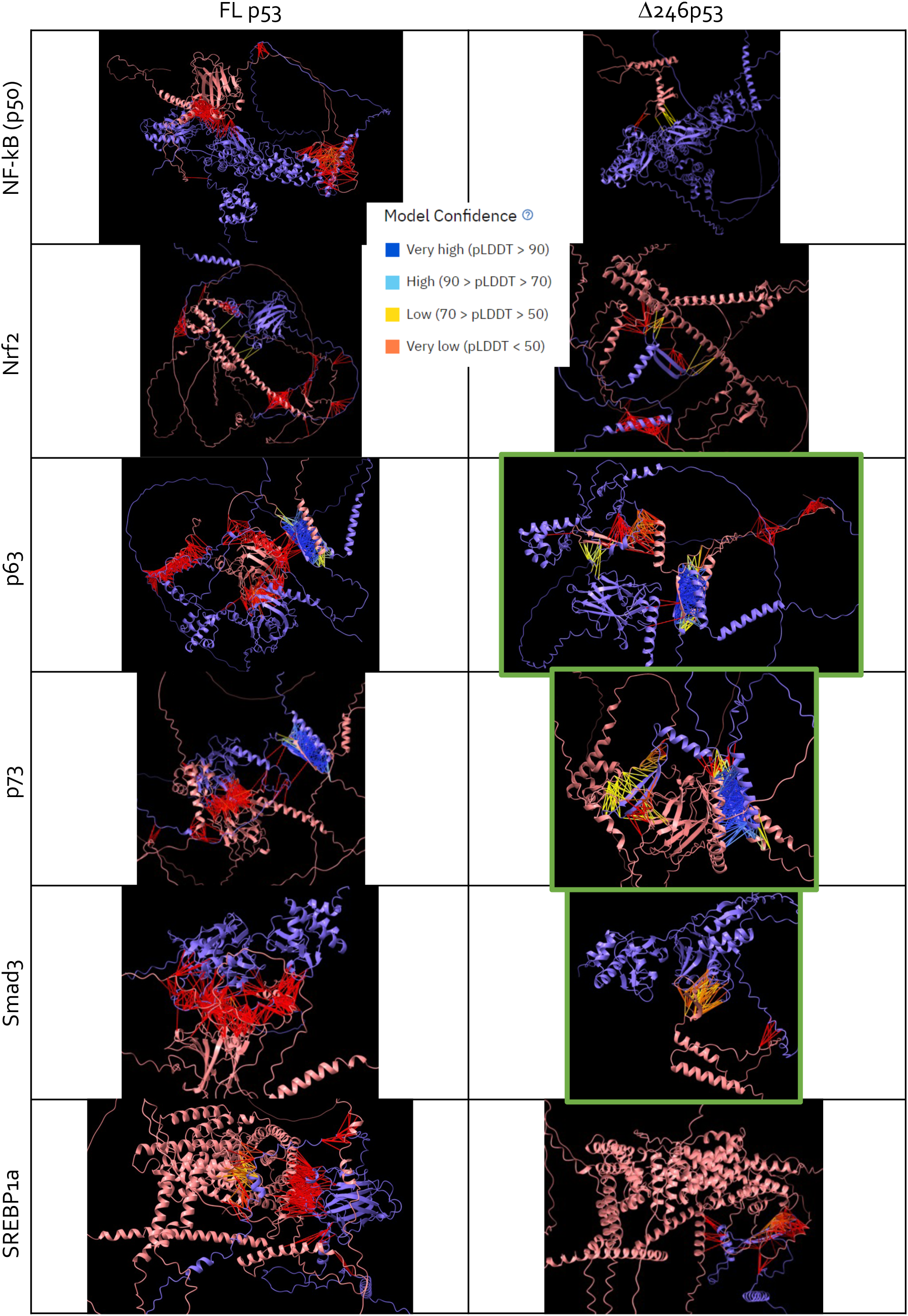
AlphaFold2-Multimer binding predictions of full-length p53 (FLp53) or Δ246p53 with candidate mutant p53-interacting proteins. The images are representative models generated by AlphaFold2-Multimer of FLp53 (left) or Δ246p53 (right) in predicted binding with already established mutant p53-interacting proteins: NF-κB(p50), NRF2, p63, p73, SMAD3 and SREBP1a. Each panel shows a predicted complex with color-coded confidence scores based on predicted Local Distance Difference Test (pLDDT) values: very high confidence (blue, pLDDT > 90), high confidence (cyan, 90 > pLDDT > 70), low confidence (yellow, 70 > pLDDT > 50), very low confidence (orange/red, pLDDT < 50). Complexes predicted with Δ246p53 highlighted in green show strongest differences as compared to their interactions with FLp53.

**Supplementary Fig. S9.**
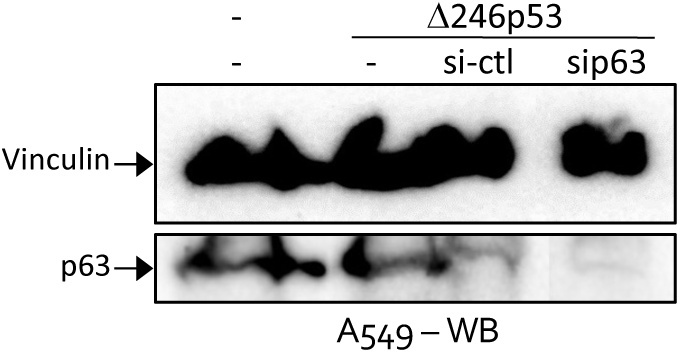
*p63* knock-down in A549 cells using siRNA. Shown is representative datum of three independent experiments.

**Supplementary Fig. S10.**
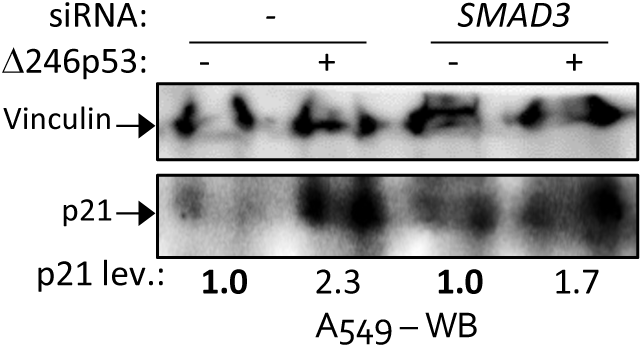
Effects of Δ246p53 and *SMAD3* in *p21* expression. Western blotting (WB) of A549 cells expressing or not Δ246p53 and treated or not with siRNA against *SMAD3* mRNA, as indicated. Shown are representative data of three independent experiments.

**Supplementary Fig. S11.**
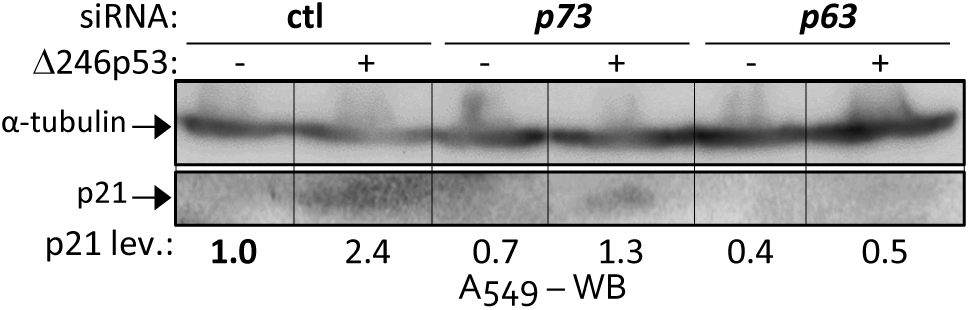
Effects of Δ246p53, p73 and p63 in *p21* expression. WB of A549 cells expressing or not Δ246p53 and treated with control siRNA (si-ctl) or siRNA against *p73* or *p63* mRNAs, as indicated. Shown is representative datum of three independent experiments.

**Supplementary Fig. S12.**
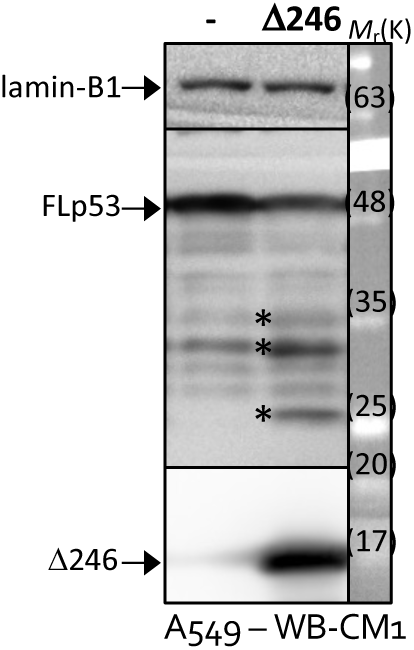
Δ246p53 expression in A549 cells. Western blotting of A549 cells expressing or not Δ246p53 construct using polyclonal anti-p53 antibody CM1. Bands marked with * are or overlap with post-translational modifications of Δ246p53, also observed in Fig. 2C.

## REFERENCES

1 Lane DP, Cheok CF, Brown C, Madhumalar A, Ghadessy FJ, Verma C. Mdm2 and p53 are highly conserved from placozoans to man. Cell Cycle 2010; 9: 540–547.

2 Candeias MM. The can and can’t dos of p53 RNA. Biochimie 2011; 93: 1962–1965.

3 Marcel V, Dichtel-Danjoy ML, Sagne C, Hafsi H, Ma D, Ortiz-Cuaran S et al. Biological functions of p53 isoforms through evolution: lessons from animal and cellular models. Cell Death Differ 2011; 18: 1815–1824.

4 Sayan BS, Sayan AE, Knight RA, Melino G, Cohen GM. p53 Is Cleaved by Caspases Generating Fragments Localizing to Mitochondria. Journal of Biological Chemistry 2006; 281: 13566–13573.

5 Campbell H, Fleming N, Roth I, Mehta S, Wiles A, Williams G et al. Δ133p53 isoform promotes tumour invasion and metastasis via interleukin-6 activation of JAK-STAT and RhoA-ROCK signalling. Nat Commun 2018; 9: 254.

6 Candeias MM, Hagiwara M, Matsuda M. Cancer-specific mutations in p53 induce the translation of Δ160p53 promoting tumorigenesis. EMBO Rep 2016; 17: 1542–1551.

7 Levine AJ. p53: 800 million years of evolution and 40 years of discovery. doi:10.1038/s41568-020-0262-1.

8 Candeias MM, Powell DJ, Roubalova E, Apcher S, Bourougaa K, Vojtesek B et al. Expression of p53 and p53/47 are controlled by alternative mechanisms of messenger RNA translation initiation. Oncogene 2006; 25: 6936–6947.

9 Fujita K, Mondal AM, Horikawa I, Nguyen GH, Kumamoto K, Sohn JJ et al. P53 Isoforms Δ133P53 and P53Β Are Endogenous Regulators of Replicative Cellular Senescence. Nat Cell Biol 2009; 11: 1135–1142.

10 Marcel V, Vijayakumar V, Fernandez-Cuesta L, Hafsi H, Sagne C, Hautefeuille A et al. p53 regulates the transcription of its Delta133p53 isoform through specific response elements contained within the TP53 P2 internal promoter. Oncogene 2010; 29: 2691–2700.

11 Gong L, Pan X, Chen H, Rao L, Zeng Y, Hang H et al. p53 isoform Δ133p53 promotes efficiency of induced pluripotent stem cells and ensures genomic integrity during reprogramming. Sci Rep 2016; 6: 37281.

12 Wang Y-H, Ho TLF, Hariharan A, Goh HC, Wong YL, Verkaik NS et al. Rapid recruitment of p53 to DNA damage sites directs DNA repair choice and integrity. Proceedings of the National Academy of Sciences 2022; 119. doi:10.1073/pnas.2113233119.

13 López-Iniesta MJ, Lacerda R, Ramalho AC, Parkar SN, Marques-Ramos A, Pereira B, et al. Internal Translation of p53 Oncoproteins During Integrated Stress Response Confers Survival Advantage on Cancer Cells. bioRxiv 2023; : 2023.03.03.531004.

14 Maier B, Gluba W, Bernier B, Turner T, Mohammad K, Guise T et al. Modulation of mammalian life span by the short isoform of p53. Genes Dev 2004; 18: 306–319.

15 Tyner SD, Venkatachalam S, Choi J, Jones S, Ghebranious N, Igelmann H et al. p53 mutant mice that display early ageing-associated phenotypes. Nature 2002; 415: 45–53.

16 Liu D, Ou L, Clemenson GD, Chao C, Lutske ME, Zambetti GP et al. Puma is required for p53-induced depletion of adult stem cells. Nat Cell Biol 2010; 12: 993–998.

17 Armata HL, Garlick DS, Sluss HK. The Ataxia Telangiectasia–Mutated Target Site Ser18 Is Required for p53-Mediated Tumor Suppression. Cancer Res 2007; 67: 11696–11703.

18 López-Iniesta MJ, Parkar SN, Ramalho AC, Lacerda R, Costa IF, Zhao J et al. Conserved Double Translation Initiation Site for Δ160p53 Protein Hints at Isoform’s Key Role in Mammalian Physiology. Int J Mol Sci 2022; 23: 15844.

19 Chen J-H, Ozanne SE, Hales CN. Methods of Cellular Senescence Induction Using Oxidative Stress. 2007, pp 179–189.

20 Mirdita M, Schütze K, Moriwaki Y, Heo L, Ovchinnikov S, Steinegger M. ColabFold: making protein folding accessible to all. Nat Methods 2022; 19: 679–682.

21 Katoh K. MAFFT: a novel method for rapid multiple sequence alignment based on fast Fourier transform. Nucleic Acids Res 2002; 30: 3059–3066.

22 Pedersen AG, Nielsen H. Neural network prediction of translation initiation sites in eukaryotes: perspectives for EST and genome analysis. Proc Int Conf Intell Syst Mol Biol 1997; 5: 226–33.

23 Vijayakumaran R, Tan KH, Miranda PJ, Haupt S, Haupt Y. Regulation of Mutant p53 Protein Expression. Front Oncol 2015; 5. doi:10.3389/fonc.2015.00284.

24 Wiley CD, Campisi J. From Ancient Pathways to Aging Cells—Connecting Metabolism and Cellular Senescence. Cell Metab 2016; 23: 1013–1021.

25 Georgakopoulou E, Evangelou K, Havaki S, Townsend P, Kanavaros P, Gorgoulis VG. Apoptosis or senescence? Which exit route do epithelial cells and fibroblasts preferentially follow? Mech Ageing Dev 2016; 156: 17–24.

26 de Larco JE, Todaro GJ. Growth factors from murine sarcoma virus-transformed cells. Proceedings of the National Academy of Sciences 1978; 75: 4001–4005.

27 Powell DJDJJ, Hrstka R, Candeias M, Bourougaa K, Vojtesek B, Fahraeus R et al. Stress-dependent changes in the properties of p53 complexes by the alternative translation product p53/47. Cell Cycle 2008; 7: 950–959.

28 Gencel-Augusto J, Lozano G. p53 tetramerization: at the center of the dominant-negative effect of mutant p53. Genes Dev 2020; 34: 1128–1146.

29 Jackson JG, Pereira-Smith OM. p53 Is Preferentially Recruited to the Promoters of Growth Arrest Genes p21 and GADD45 during Replicative Senescence of Normal Human Fibroblasts. Cancer Res 2006; 66: 8356–8360.

30 Lahav G, Rosenfeld N, Sigal A, Geva-Zatorsky N, Levine AJ, Elowitz MB et al. Dynamics of the p53-Mdm2 feedback loop in individual cells. Nat Genet 2004; 36: 147–150.

31 Candeias MM, Powell DJ, Roubalova E, Apcher S, Bourougaa K, Vojtesek B et al. Expression of p53 and p53/47 are controlled by alternative mechanisms of messenger RNA translation initiation. Oncogene 2006; 25: 6936–6947.

32. 32 Candeias MM. Mutant p53 loses “gain-of-functions” after losing isoform expression. In: 6th International Mutant p53 Workshop. Cell Death and Differentiation Conferences: Toronto, 2013, p 1.

33 Yin Y, Stephen CW, Luciani MG, Fahraeus R. p53 Stability and activity is regulated by Mdm2-mediated induction of alternative p53 translation products. Nat Cell Biol 2002; 4: 462–467.

34 McLure KG. How p53 binds DNA as a tetramer. EMBO J 1998; 17: 3342–3350.

35 Keyes WM, Mills AA. p63: A New Link Between Senescence and Aging. Cell Cycle 2006; 5: 260–265.

36 Guo X, Keyes WM, Papazoglu C, Zuber J, Li W, Lowe SW et al. TAp63 induces senescence and suppresses tumorigenesis in vivo. Nature Cell Biology 2009 11:12 2009; **11**: 1451–1457.

37 Westfall MD, Mays DJ, Sniezek JC, Pietenpol JA. The Delta Np63 alpha phosphoprotein binds the p21 and 14-3-3 sigma promoters in vivo and has transcriptional repressor activity that is reduced by Hay-Wells syndrome-derived mutations. Mol Cell Biol 2003; 23: 2264–2276.

38 Petitjean A, Ruptier C, Tribollet V, Hautefeuille A, Chardon F, Cavard C et al. Properties of the six isoforms of p63: p53-like regulation in response to genotoxic stress and cross talk with ΔNp73. Carcinogenesis 2008; 29: 273–281.

39 Moore L, Lu X, Ghebranious N, Tyner S, Donehower LA. Aging-associated truncated form of p53 interacts with wild-type p53 and alters p53 stability, localization, and activity. Mech Ageing Dev 2007; 128: 717–730.

40 Nakagawa S, Niimura Y, Gojobori T, Tanaka H, Miura K -i. Diversity of preferred nucleotide sequences around the translation initiation codon in eukaryote genomes. Nucleic Acids Res 2007; 36: 861–871.

41 López-Iniesta MJ, Lacerda R, Ramalho AC, Parkar SN, Marques-Ramos A, Pereira B, et al. Internal Translation of p53 Oncoproteins During Integrated Stress Response Confers Survival Advantage on Cancer Cells. bioRxiv 2023; : 2023.03.03.531004.

42 López-Iniesta MJ, Parkar SN, Ramalho AC, Lacerda R, Costa IF, Zhao J et al. Conserved Double Translation Initiation Site for Δ160p53 Protein Hints at Isoform’s Key Role in Mammalian Physiology. Int J Mol Sci 2022; 23: 15844.

43 Cordani M, Oppici E, Dando I, Butturini E, Dalla Pozza E, Nadal-Serrano M et al. Mutant p53 proteins counteract autophagic mechanism sensitizing cancer cells to mTOR inhibition. Mol Oncol 2016; 10: 1008–1029.

44 Lisek K, Campaner E, Ciani Y, Walerych D, Del Sal G. Mutant p53 tunes the NRF2-dependent antioxidant response to support survival of cancer cells. Oncotarget 2018; 9: 20508–20523.

45 Gaiddon C, Lokshin M, Ahn J, Zhang T, Prives C. A Subset of Tumor-Derived Mutant Forms of p53 Down-Regulate p63 and p73 through a Direct Interaction with the p53 Core Domain. Mol Cell Biol 2001; 21: 1874.

46 Adorno M, Cordenonsi M, Montagner M, Dupont S, Wong C, Hann B et al. A Mutant-p53/Smad Complex Opposes p63 to Empower TGFβ-Induced Metastasis. Cell 2009; 137: 87–98.

47 Freed-Pastor WA, Mizuno H, Zhao X, Langerod A, Moon SH, Rodriguez-Barrueco R et al. Mutant p53 disrupts mammary tissue architecture via the mevalonate pathway. Cell 2012; 148: 244–258.

